# The phylogeography of westslope cutthroat trout

**DOI:** 10.1101/213363

**Authors:** Michael K. Young, Kevin S. McKelvey, Tara Jennings, Katie Carter, Richard Cronn, Ernest R. Keeley, Janet L. Loxterman, Kristy L. Pilgrim, Michael K. Schwartz

## Abstract

Identifying units of conservation of aquatic species is fundamental to informed natural resources science and management. We used a combination of mitochondrial and nuclear molecular methods to identify potential units of conservation of westslope cutthroat trout, a taxon native to montane river basins of the northwestern U.S. and southwestern Canada. Mitogenomic sequencing identified two major lineages composed of nine monophyletic clades, and a well-supported subclade within one of these, largely delineated by river basins. Analyses of microsatellites and single nucleotide polymorphisms corroborated most of these groupings, sometimes with less resolution but demonstrating more complex connections among clades. The mitochondrial and nuclear analyses revealed that Pleistocene glacial cycles profoundly influenced the distribution and divergence of westslope cutthroat trout, that this taxon crossed the Continental Divide in two separate events, and that genetically pure but nonindigenous fish were widely distributed. Herein, we recognize nine geographically discrete, cytonuclear lineages largely circumscribed by major river basins as potential units of conservation: 1) John Day; 2) Coeur d’Alene; 3) St. Joe; 4) North Fork Clearwater; 5) Salmon; 6) Clearwater headwaters; 7) Clearwater-eastern Cascades; 8) neoboreal, consisting of most of the Columbia upstream from central Washington, the Fraser in British Columbia, and the South Saskatchewan in Alberta; and 9) Missouri.

---

> If the biota, in the course of aeons, has built something we like but do not understand, then who but a fool would discard seemingly useless parts? To keep every cog and wheel is the first precaution of intelligent tinkering.
>
> —Aldo Leopold (1953)

Leopold’s (1953) admonition has long served as a core paradigm of conservation biology. In his time, the focus was on preserving species, but species often comprise distinct lineages that go by many names—subspecies, distinct population segments, evolutionarily significant units, operational taxonomic units, or stocks—and constitute potential units of conservation (Avise 2000) that can be accorded taxonomic or legal standing, e.g., under the U.S. Endangered Species Act (Waples 1991) or Canada’s Species at Risk Act (Mee et al. 2015). Historically, evidence for the presence of these lineages was largely based on variation in morphology, behavior, life history, or ecology, usually associated with a geographically discrete distribution, but increasingly sophisticated molecular tools have revolutionized their identification and spawned a branch of science dedicated to their study, phylogeography (Avise 2000). There is particular urgency to identify units of conservation for aquatic ectotherms such as fishes, mussels, crayfish, and amphibians. These aquatic groups are disproportionately represented among at-risk taxa (Williams et al. 2011), and shifts in climate are beginning to alter their distributions (Comte and Grenouillet 2013; Eby et al. 2014). Correctly inferring the existence and extent of lineages within a taxon also relies on an understanding of the geological and climatic histories of current and former habitats. For aquatic species, especially fishes in western North America, this means appreciating the variation in hydrological connections over evolutionary timescales (McPhail and Lindsey 1986; Minckley et al. 1986), particularly with respect to the influence of Pleistocene glaciations on hydrologic networks (Bernatchez and Wilson 1998). A robust intraspecific phylogeny also demands that a taxon be sampled across its entire range with sufficient intensity to represent all potential units of conservation, and be examined with tools of sufficient sensitivity to recognize them (Heath et al. 2008). It also requires awareness of the potential for human-assisted relocation of nonlocal fish populations (Metcalf et al. 2012; Pritchard et al. 2015).

The systematics and distribution of cutthroat trout (*Oncorhynchus clarkii*) have been a source of contention since the species was first formally described in the 19^th^ century (Richardson 1836). Various forms have been described as full species or as subspecies with little regard to previous work (Thorgaard et al., this volume), and the origin and affiliation of some type specimens has been obscure (Metcalf et al. 2012). Moreover, our understanding of the historical distributions of putative taxa is far from perfect, in part because of the broad-scale husbandry and translocation of some forms of cutthroat trout (Wiltzius 1985) and the rapid extirpation of others (Young and Harig 2001). A valiant attempt to resolve this ambiguity was made by the preeminent taxonomist of the 20^th^ century on western North American trout, Robert Behnke, who divided cutthroat trout into 14 extant and 2 extinct subspecies (Behnke 1992). Among these subspecies was the westslope cutthroat trout *O. c. lewisi*, first observed by Lewis and Clark in 1805 at the eventual type location, the Great Falls of the Missouri River (Girard 1856). Primarily relying on morphological evidence, Behnke (1992) asserted that the cutthroat trout of the interior Columbia River basin also represented this subspecies, an observation broadly corroborated by phylogenetic work (Leary et al. 1987; Wilson and Turner 2009; Loxterman and Keeley 2012; Houston et al. 2012). Whether there is substantial phylogenetic diversity within this taxon across its range, however, has not been resolved. Because of the absence of substantial morphological differences among westslope cutthroat trout in different river basins, the U.S. Fish and Wildlife Service concluded that the taxon constituted a single unit of conservation in response to a petition for listing under the U.S. Endangered Species Act (U.S. Fish and Wildlife Service 2003). Yet subsequent molecular studies indicated the presence of marked geographic divergence within westslope cutthroat trout (Drinan et al. 2011; Loxterman and Keeley 2012) that was likely driven by the complex geological and climatic history of the region (McPhail and Lindsey 1986).

Although Behnke (1992) was uncertain about the ultimate source of the lineage that led to westslope cutthroat trout, he posited that most of its present distribution could be attributed to events during the Wisconsinan glaciation 100–15 ka ago. He proposed that major waterfalls developed during this time in the Pend Oreille, Spokane, and Kootenai River basins that may have isolated westslope cutthroat trout upstream. Thereafter, the Clark Fork River basin served as the source for fish dispersing—across divides via stream capture—to the east and north to the upper Missouri River and South Saskatchewan River basins in Montana, Wyoming, and Alberta and to the south to the Clearwater and Salmon River basins in Idaho. He postulated that outwash floods from Glacial Lake Missoula carried westslope cutthroat trout to disjunct river basins along the eastern face of the Cascade Range in Washington and to the John Day River basin in Oregon, and may have been an alternate route for populations that colonized the Clearwater and Salmon River basins. Recession of continental glaciers, along with the temporary proglacial lakes at their trailing edges, presumably permitted westslope cutthroat trout to colonize waters farther north in western Montana, northern Idaho, southern British Columbia, and southern Alberta.

Our goal was to test this phylogeographic hypothesis (cf. Crisp et al. 2011). We did so by assessing the phylogenetic structure of westslope cutthroat trout across the bulk of its historical distribution, with an additional aim to identify and geographically delineate potential units of conservation, using a combination of mitochondrial and nuclear molecular information on different samples of westslope cutthroat trout, including those used to establish hatchery broodstocks.

## Methods

### The historical range and its geological history

The historical range of westslope cutthroat trout is thought to constitute portions of the Fraser, South Saskatchewan, Missouri, and Columbia River basins (Shepard et al. 2005; McPhail 2007; Figure 1). Its presence in the Fraser River basin in British Columbia is limited to a few headwater streams (McPhail 2007), whereas it once occupied most of the tributaries in the Bow River and Oldman River basins, part of the South Saskatchewan River basin in Alberta (Yau and Taylor 2013; Fisheries and Oceans Canada 2014). In the Missouri River basin, its range was thought to include the main stem and most tributaries from the Judith River to the headwaters, excluding those blocked by waterfalls (e.g., the Sun River basin above present-day Gibson Dam). In the upper Columbia River basin, the historical range includes 1) the entire Pend Oreille and Kootenai River basins in British Columbia, Idaho, Montana, and Washington; 2) scattered populations in the Kettle River basin in British Columbia (McPhail 2007) and presumably Washington; 3) the Spokane River basin above Spokane Falls, primarily in Idaho, and perhaps the Sanpoil River basin in Washington (Trotter et al. 2001b). Along the eastern Cascades, westslope cutthroat trout are found from the Methow River basin in the north to the Yakima River basin in the south, although there is uncertainty with respect to whether it is indigenous to these and all intervening basins (Trotter et al. 1999, 2001a). In Oregon, westslope cutthroat trout are present in tributaries of the main-stem and North Fork John Day River, but believed to be introduced to the latter (Gunckel 2002). Westslope cutthroat trout are also native to the bulk of two large river basins in the Snake River in Idaho, the Clearwater and Salmon, but are curiously absent from all other basins tributary to the Snake River in Oregon, Washington, and Idaho (except where introduced; Neville and Dunham 2011). Although the four major basins are presently hydrologically isolated, they were presumably connected in the distant or recent past. For example, the upper Columbia and Missouri Rivers were connected by the dual outlets of a lake at Marias Pass in Montana until 1913, when the Union Pacific Railroad diverted all flow to a tributary of the Two Medicine River of the Missouri River basin (Schultz 1941).

**Figure 1.**
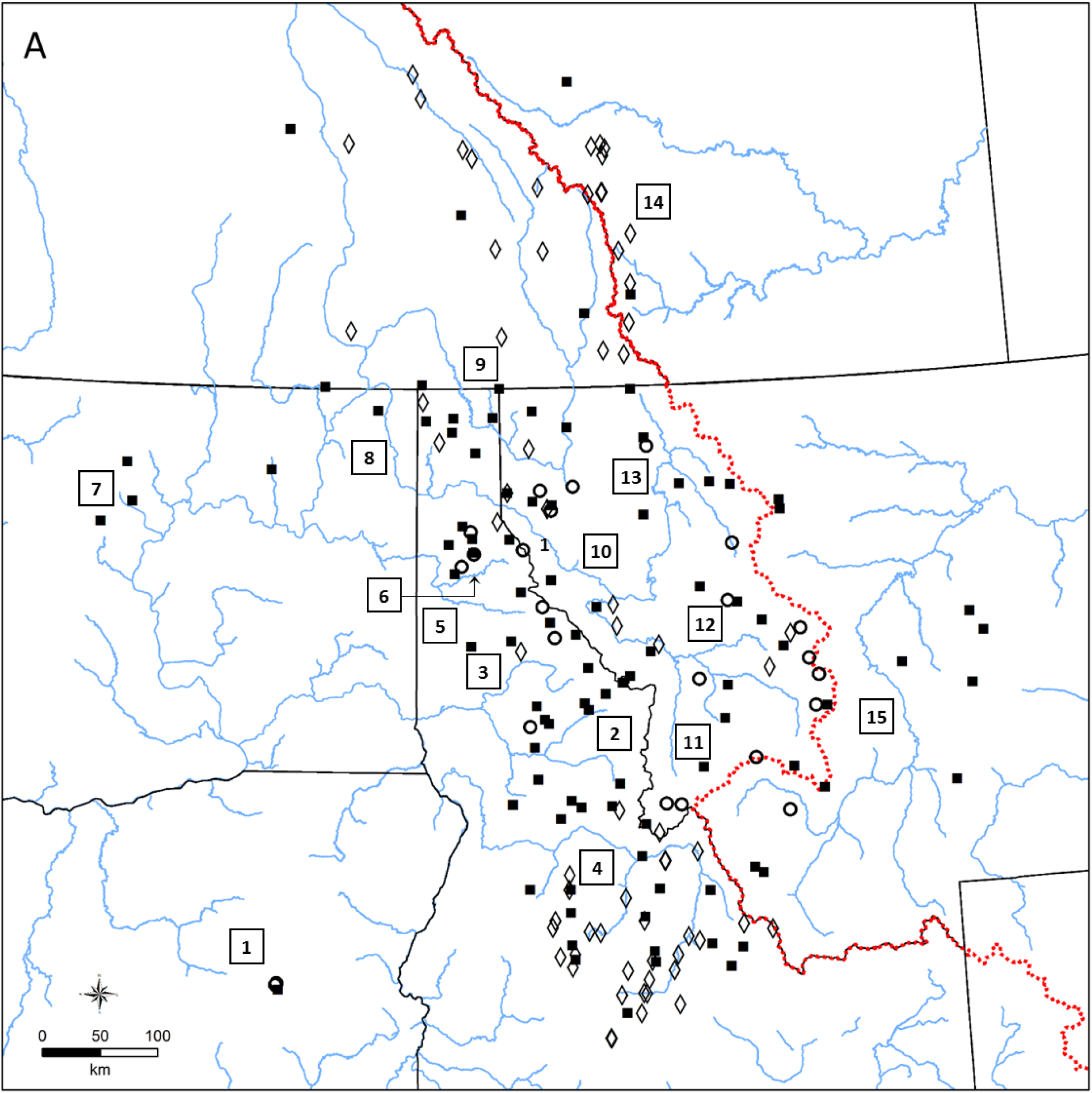
Sample locations, river basins, and units of conservation within the historical range of westslope cutthroat trout. (A) Sources of westslope cutthroat trout specimens that were genetically analyzed using mitogenomic sequences and microsatellites (squares), ND2 sequences (diamonds), or SNPs (circles). Fish from some locations were analyzed using more than one method. Major river basins are: 1, John Day; 2, Clearwater; 3, North Fork Clearwater; 4, Salmon; 5, St. Joe; 6, Coeur d’Alene; 7, Wenatchee River, Lake Chelan, and Methow River in the eastern Cascades; 8, Pend Oreille; 9, Kootenai; 10, Clark Fork; 11, Bitterroot; 12, Blackfoot; 13, Flathead; 14, Bow (north) and Oldman (south) Rivers in the South Saskatchewan; and 15, Missouri. (B) Extent of proposed units of conservation. Colored polygons correspond to those identifying mitogenomic clades (Salmon River clades are combined). The dotted red line denotes the Continental Divide.

The modern distribution has been substantially reduced because of habitat alteration and introductions of nonnative species (Shepard et al. 2005). Moreover, hatchery strains of westslope cutthroat trout have been widely introduced throughout the 20^th^ century and could affect both the phylogenetic interpretation and evolutionary integrity of wild populations. Three broodstocks are commonly used at present. In Montana, the primary broodstock was developed from fish in the upper Flathead and middle Clark Fork River basins (Drinan 2010). In Idaho, westslope cutthroat trout were domesticated from populations in the Priest River basin, as was the King's Lake broodstock in Washington (Crawford 1979; M. Campbell, Idaho Department of Fish and Game, personal communication). A second broodstock in Washington, Twin Lakes, was presumably developed from fish collected in the Lake Chelan basin or Methow or Wenatchee River basins (Crawford 1979).

The interior Columbia River basin constitutes the western portion of the distribution of westslope cutthroat trout and has been present in some form for at least 15 Ma, roughly the age of the lineage representing cutthroat trout and rainbow trout *O. mykiss* (Stearley and Smith 2016). The eruption and deposition of flood basalts from the mid-Miocene to the Pliocene led to repeated re-routing and re-entrenchment of the major tributaries (Fecht et al. 1985). This coincided with the orogeny of the Cascade Range, whereas the topography of the headwaters was set by the orogeny of the Rocky Mountains 60 Ma earlier (Alt and Hyndman 1995). The Pleistocene was characterized by the cyclic advance and retreat of the Cordilleran and Laurentide ice sheets, which buried northerly portions of the current range of westslope cutthroat trout under hundreds of meters of ice. The repeated advance of the Purcell lobe of the Cordilleran ice sheet near Sandpoint, Idaho also led to the damming of the Clark Fork River and filling of Glacial Lake Missoula (Waitt 1985). The repeated failure of this ice dam led to catastrophic jökulhlaups lasting 1–2 weeks in which the discharge from the draining glacial lake temporarily exceeded that of all rivers on Earth combined. These glacial outburst floods, perhaps up to 100 at multi-decadal intervals during the Wisconsinan glaciation (Booth et al. 2003) and an unknown number from earlier Pleistocene glacial intervals (Bader et al. 2016), produced the Columbia Basin scablands. The scablands are characterized by floodwater-scoured surfaces, deep sedimentary deposits from temporary ponding as water backed up at topographic chokepoints, e.g., Wallula Gap in Washington, and huge nickpoints in many river channels that resulted in waterfalls likely to constitute migration barriers to aquatic taxa, e.g., Spokane Falls on the Spokane River, Palouse Falls on the Palouse River, Metalline Falls on the Pend Oreille River, and falls on many lesser basins (Waitt 1985). The Okanagan and Columbia River lobes of this ice sheet also led to the formation of Glacial Lake Columbia, which at its highstand extended upstream to the Purcell lobe and portions of the Coeur d’Alene and St. Joe River valleys (Atwater 1987; Hanson and Clague 2016). Contemporaneous mountain glaciers (Pierce 2003) would have forced headwater taxa to lower portions of many watersheds to escape habitats that were cold, turbid, and unproductive. Similar conditions—rerouted and glacially derived streams and temporary glacial lakes—prevailed in the upper Missouri River basin. Until the Pleistocene (and perhaps during portions of it), the Missouri River drained northward to Hudson Bay (Howard 1958). By the Last Glacial Maximum 23–19 ka ago (Hughes et al. 2013), the Laurentide ice sheet had forced the Missouri River south to near its present course within the Mississippi River basin. As that ice sheet retreated, it created a series of proglacial lakes (Alden 1932; Colton et al. 1961) that persisted as recently as 11.5 ka ago (Davis et al. 2006) and may have served as springboards for colonization by aquatic taxa farther north. The retreating Cordilleran ice sheet also left a trail of temporary lakes that could have served as stepping stones for aquatic species to colonize basins farther north, such as the Fraser River (McPhail and Lindsey 1986).

Two other major events in the history of the Columbia River basin merit attention. About 3 Ma ago, the Columbia River captured the Snake River, which previously drained farther south to the Klamath or Sacramento River basins (Wood and Clemens 2002; Stearley and Smith 2016). Although the connection between the Snake River and lower Columbia River did not promote extensive faunal transfer (at least of salmonids), it did provide a route for another Pleistocene flood: the draining of Lake Bonneville. Unlike the ice dam failures of Glacial Lake Missoula, Lake Bonneville drained by overtopping and rapidly downcutting at Red Rock Pass 17.5 ka ago (Amidon and Clark 2015; Oviatt 2015), delivering 4,750 km^3^ of water in what represents the largest (by volume) known freshwater flood (O’Connor 1993). Like those of Glacial Lake Missoula, this flood backed up many of the tributary rivers and may have conveyed aquatic species across large distances. Because it was less violent and longer lasting—a couple months to one year (Malde 1968; Jarrett and Malde 1987)—the draining of Lake Bonneville may been more conducive to fish survival and translocation. Interpreting its possible phylogeographic effect is difficult because it was both preceded and followed by outwash floods from Glacial Lake Missoula, the last of which was 14.7 ka ago (Waitt 1985; McDonald et al. 2012; Balbas et al. 2017), with additional floods still later from smaller proglacial lakes that formed as the Purcell lobe of the Cordilleran ice sheet retreated (Waitt et al. 2009; Peters 2012) and perhaps from the longer-persisting Glacial Lake Columbia (Balbas et al. 2017).

### Conservation units

We define units of conservation (sensu distinct population segments; Waples 1991) as populations or groups thereof that are substantially reproductively isolated and that constitute an important component of the evolutionary legacy of a species. Ideally, units of conservation would be identified based on an array of phylogenetically informative ecological, life history, morphological, and genetic characteristics (Fraser and Bernatchez 2001). Variation in morphology has been related to ecotypic differences in westslope cutthroat trout (Seiler and Keeley 2009), and such differences have been shown to have a genetic basis and contribute to identifying units of conservation in some salmonids (Keeley et al. 2007). To our knowledge, however, there has been no comparison of morphological or ecological traits across the entire range of westslope cutthroat trout (cf. Bestgen et al. 2013). In addition, relative to other Pacific salmon and trout, cutthroat trout are ecological generalists capable of expressing a wide array of life history strategies (Young 1995; Waples et al. 2001), rendering these characteristics less definitive for delineating units of conservation. Given this background of uncertainty, we used well-supported, reciprocally monophyletic mitochondrial clades (Moritz et al. 1995) as working hypotheses about the units of conservation within westslope cutthroat trout, particularly when geographic structuring reinforced those distinctions. We also sought support from analyses of two suites of nuclear markers to corroborate the mitochondrial results.

### Laboratory analyses

We obtained three sets of genetic information from two groups of samples of westslope cutthroat trout: 1) mitogenomic sequences of specimens from most of the historical range in the U.S. and Canada (*n* = 96 fish); 2) allele frequencies of 10 microsatellite loci (Vu and Kalinowski 2009) in the aforementioned specimens (save one from Buck Creek, WA, sample 44); and 3) allele frequencies of 52 variable nuclear single nucleotide polymorphisms (SNPs) from specimens collected across a large portion of the U.S. range (*n* = 524 fish from 55 sites; Table 1; Figure 1). Specimens from wild populations that were representative of those used to establish each of the hatchery broodstocks were included in one or both sample groups. Some of the sites in the first two analyses also had additional individuals analyzed using SNPs (*n* = 29), but spatial overlap in these datasets was not complete.

**Table 1.** Samples used in the phylogeographic analysis of westslope cutthroat trout. Site labels are in Figure 1.

| Site | 4th-code HU<sup>1</sup> | State/<br>Province | Latitude | Longitude | N<sup>2</sup> | MG<sup>3</sup> | ND2<sup>4</sup> | SNP<sup>5</sup> | Label |
| --- | --- | --- | --- | --- | --- | --- | --- | --- | --- |
| Albert | Middle Clark Fork | MT | 46.9213 | -114.3678 | 1 | 1 | A |  | 1 |
| Beaver | Yaak | MT | 48.8174 | -115.6787 | 1 | 2 | A |  | 2 |
| Bostwick | Gallatin | MT | 45.7995 | -110.9981 | 10 | 3 | A | x | 3 |
| Moose | Blackfoot | MT | 46.9227 | -112.8534 | 6 | **4** | A | x | 4 |
| Deer | Bitterroot | MT | 45.5746 | -114.4784 | 13 | 5 | A | x | 5 |
| EF Bull | Lower Clark Fork | MT | 48.1151 | -115.6868 | 11 | 6 | B | x | 6 |
| Ketchikan | North Fork Flathead | MT | 48.9771 | -114.5071 | 1 | 7 | A |  | 7 |
| McGuire | Upper Kootenai | MT | 48.6896 | -115.2728 | 10 | 8 | A | x | 8 |
| Martin | Bitterroot | MT | 46.0057 | -113.8161 | 1 | 9 | A |  | 9 |
| McCabe | Blackfoot | MT | 47.1324 | -113.0840 | 6 | 10 | A | x | 10 |
| Miller | Fisher | MT | 48.0827 | -115.4642 | 5 | 11 | A | x | 11 |
| Meadow | Lower Selway | ID | 45.7154 | -115.1953 | 12 | 12 | C | x | 12 |
| Norton | Upper Clark Fork | MT | 45.9811 | -112.8041 | 8 | 13 | A | x | 13 |
| Ontario | Upper Clark Fork | MT | 46.4454 | -112.3940 | 8 | 14 | A | x | 14 |
| North | Bitterroot | MT | 46.7343 | -114.6085 | 1 | 15 | A |  | 15 |
| Tyler | Flint-Rock | MT | 46.6377 | -113.5047 | 1 | 16 | A |  | 16 |
| Wilkes | Lower Clark Fork | MT | 47.4962 | -115.4883 | 1 | 17 | A |  | 17 |
| Canuck | Moyie | MT | 48.9988 | -116.0618 | 8 | 18 | A | x | 18 |
| Ball | Lower Kootenai | ID | 48.7712 | -116.6069 | 10 | 19 | A | x | 19 |
| Rampike | Upper Coeur d'Alene | ID | 47.8197 | -115.9619 | 10 | 20 | D | x | 20 |
| Trail | Upper Coeur d'Alene | ID | 47.8291 | -116.3941 | 10 | 21 | D | x | 21 |
| West Gold | Pend Oreille Lake | ID | 47.9259 | -116.5078 | 10 | 22 | A | x | 22 |
| Avalanche | Upper Missouri | MT | 46.7439 | -111.5180 | 7 | **23** | A | x | 23 |
| Beaver | Priest | ID | 48.7489 | -116.9270 | 10 | 24 | A | x | 24 |
| Mokins | Upper Spokane | ID | 47.7826 | -116.6663 | 1 | 25 | A |  | 25 |

| Site | 4th-code HU<sup>1</sup> | State/<br>Province | Latitude | Longitude | *N*<sup>2</sup> | MG<sup>3</sup> | ND2<sup>4</sup> | SNP<sup>5</sup> | Label |
| --- | --- | --- | --- | --- | --- | --- | --- | --- | --- |
| NF St. Joe | St. Joe | ID | 47.4055 | -115.8382 | 1 | 26 | B |  | 26 |
| Skin | Moyie | MT | 48.7728 | -116.1446 | 1 | 27 | A |  | 27 |
| Slowey | Middle Clark Fork | MT | 47.2808 | -114.9731 | 1 | 28 | A |  | 28 |
| Dry Wolf | Judith | MT | 46.9492 | -110.5684 | 1 | 29 | A |  | 29 |
| Fish | Jefferson | MT | 45.8032 | -112.4725 | 1 | **30** | A |  | 30 |
| Heller | St. Joe | ID | 47.0662 | -115.2167 | 1 | 31 | E |  | 31 |
| Tributary of Armstrong | Coeur d'Alene Lake | ID | 47.5543 | -116.5993 | 1 | 32 | A |  | 32 |
| NF Dupuyer | Two Medicine | MT | 48.0636 | -112.8182 | 1 | 33 | A |  | 33 |
| Sawmill | Belt | MT | 47.1013 | -110.7097 | 1 | **30** | A |  | 34 |
| Youngs | South Fork Flathead | MT | 47.2838 | -113.3568 | 1 | 35 | A |  | 35 |
| French | Upper North Fork Clearwater | ID | 46.5143 | -115.6768 | 1 | **36** | C |  | 36 |
| Osier | Upper North Fork Clearwater | ID | 46.8055 | -115.0836 | 1 | 37 | F |  | 37 |
| Crooked Fork | Lochsa | ID | 46.6843 | -114.6958 | 1 | 38 | C |  | 38 |
| Pete King | Lochsa | ID | 46.1929 | -115.7064 | 1 | 39 | C |  | 39 |
| Baldy | South Fork Clearwater | ID | 45.9415 | -115.6750 | 1 | 40 | C |  | 40 |
| Bad Luck | Upper Selway | ID | 45.8956 | -114.7553 | 1 | 41 | B |  | 41 |
| Flat | Upper Selway | ID | 45.7213 | -114.8575 | 1 | 42 | C |  | 42 |
| Twisp | Methow | WA | 48.3839 | -120.4342 | 5 | 43 | G | x | 43 |
| Buck | Wenatchee | WA | 47.9163 | -120.7132 | 1 | **36** | C |  | 44 |
| Cedar | Pend Oreille | ID | 48.8333 | -117.4952 | 9 | **45** | A | x | 45 |
| Ninemile | Sanpoil | WA | 48.3596 | -118.7420 | 1 | 46 | A |  | 46 |
| Mill | Upper Salmon | ID | 44.4940 | -114.4200 | 10 | **47** | F | x | 47 |
| Morse | Pashimeroi | ID | 44.6222 | -113.7999 | 9 | **47** | F | x | 48 |
| Jacobs Ladder | Lower Middle Fork Salmon | ID | 45.0766 | -115.3373 | 10 | 49 | F | x | 49 |
| MF Little Timber | Lemhi | ID | 44.5867 | -113.4618 | 1 | 50 | F |  | 50 |
| American | South Fork Clearwater | ID | 45.7452 | -115.9635 | 1 | 51 | C |  | 51 |
| Scotchman | Flint-Rock | MT | 46.3818 | -113.5519 | 1 | 52 | H |  | 52 |
| Fourmile | Smith | MT | 46.5447 | -110.7372 | 8 | 53 | A | x | 53 |
| Kraft | Swan | MT | 47.4142 | -113.7778 | 10 | **4** | A | x | 54 |
| Leiberg | Upper Coeur d'Alene | ID | 47.7209 | -116.3834 | 10 | 55 | I | x | 55 |
| NF Grouse | Pend Oreille Lake | ID | 48.4962 | -116.3481 | 1 | 56 | J |  | 56 |
| NF Teton | Teton | MT | 47.9905 | -112.8143 | 1 | **23** | A |  | 57 |
| SF Red | South Fork Clearwater | ID | 45.6282 | -115.4310 | 10 | 58 | A | x | 58 |
| Yoosa | Clearwater | ID | 46.4100 | -115.5864 | 7 | **45** | A | x | 59 |
| Falls | Lake Chelan | WA | 48.0810 | -120.3577 | 6 | 60 | C | x | 60 |
| Thayer | Big Hole | MT | 45.2054 | -113.2944 | 1 | **61** | A |  | 61 |
| Hungery | Lochsa | ID | 46.3758 | -115.5400 | 1 | 62 | C |  | 62 |
| Ross | Upper Kootenai | MT | 48.1853 | -115.9790 | 1 | 63 | K |  | 63 |
| Ditch | South Fork Clearwater | ID | 45.7712 | -115.3051 | 10 | 64 | L | x | 64 |
| EF Emerald | St. Joe | ID | 46.9871 | -116.4163 | 1 | 65 | B |  | 65 |
| Big | Pashimeroi | ID | 44.4416 | -113.5987 | 1 | 66 | F |  | 66 |
| Bitter | Kettle HW | BC | 49.0153 | -118.1243 | 1 | 67 | B |  | 67 |
| Blackbird | Middle Salmon-Panther | ID | 45.0675 | -114.3487 | 1 | 68 | F |  | 68 |
| Blairmore | Oldman | AB | 49.7143 | -114.4630 | 1 | 69 | M |  | 69 |
| Bluff | St. Joe | ID | 47.1668 | -115.5088 | 1 | 70 | E |  | 70 |
| Boundary | Upper Middle Fork Salmon | ID | 44.5276 | -115.3004 | 1 | 71 | F |  | 71 |
| Buffalo | Beaverhead | MT | 45.1646 | -113.2000 | 1 | **61** | A |  | 72 |
| Cayuse | Upper North Fork Clearwater | ID | 46.5998 | -114.8931 | 1 | 73 | C |  | 73 |
| Colson | Middle Salmon-Panther | ID | 45.3247 | -114.5365 | 1 | 74 | N |  | 74 |
| Crazy | Fraser HW | BC | 51.0228 | -118.6053 | 1 | 75 | A |  | 75 |
| Duck | Lower Middle Fork Salmon | ID | 44.8522 | -114.5220 | 1 | 76 | F |  | 76 |
| Floodwood | Lower North Fork Clearwater | ID | 47.0241 | -115.9553 | 1 | 77 | F |  | 77 |
| Hartley | Kootenai HW | BC | 49.5761 | -115.0255 | 1 | 78 | A |  | 78 |
| Indian Grave | Lochsa | ID | 46.4787 | -115.0899 | 1 | 79 | C |  | 79 |
| John Day | Upper John Day | OR | 44.2978 | -118.5501 | 1 | 80 | O |  | 80 |
| Meadow | South Fork Salmon | ID | 44.8950 | -115.3404 | 1 | 81 | P |  | 81 |
| Monk | Priest HW | BC | 49.0329 | -116.9780 | 1 | 82 | A |  | 82 |
| NF Elkhorn | Upper Middle Fork Salmon | ID | 44.6405 | -115.3324 | 1 | 83 | Q |  | 83 |
| Pack | Pend Oreille Lake | ID | 48.6592 | -116.6244 | 1 | 84 | B |  | 84 |
| Sheep | Middle Fork Flathead | MT | 48.2321 | -113.6183 | 1 | 85 | A |  | 85 |
| Split | South Fork Salmon | ID | 45.0809 | -115.7844 | 1 | 86 | F |  | 86 |
| Toby | Columbia HW | BC | 50.3598 | -116.5000 | 1 | 87 | A |  | 87 |
| Truman | Flathead Lake | MT | 47.9938 | -114.4016 | 1 | 88 | R |  | 88 |
| Twin | Upper Salmon | ID | 44.5792 | -114.4275 | 1 | 89 | B |  | 89 |
| Waiparous | Bow | AB | 51.3864 | -115.1686 | 1 | 90 | A |  | 90 |
| Warm Springs | Upper Salmon | ID | 44.1029 | -114.7472 | 1 | 91 | F |  | 91 |
| Werner | North Fork Flathead | MT | 48.5931 | -114.3679 | 1 | 92 | S |  | 92 |
| Withington | Lemhi | ID | 45.0422 | -113.7966 | 1 | 93 | F |  | 93 |
| Wounded Buck | South Fork Flathead | MT | 48.2280 | -113.9705 | 1 | 94 | A |  | 94 |
| Gravey | Upper North Fork Clearwater | ID | 46.5284 | -115.1336 | 1 | 95 | C |  | 95 |
| Twentyfivemile | Middle Fork Flathead | MT | 48.2055 | -113.3799 | 1 | 96 | A |  | 96 |
| WF Bitterroot | Bitterroot | MT | 45.5040 | -114.3334 | 1 |  | Z |  | 97 |
| Beaver | Blackfoot | MT | 47.0190 | -112.7674 | 1 |  | Z |  | 98 |
| Chimney | Blackfoot | MT | 46.7626 | -113.0219 | 1 |  | Z |  | 99 |
| Canyon | Bow | AB | 50.8953 | -114.7774 | 1 |  | Z |  | 100 |
| Picklejar 2 | Bow | AB | 50.5198 | -114.7737 | 1 |  | Z |  | 101 |
| Picklejar 4 | Bow | AB | 50.5188 | -114.7841 | 1 |  | Z |  | 102 |
| Prairie | Bow | AB | 50.8783 | -114.8885 | 1 |  | W |  | 103 |
| Quirk | Bow | AB | 50.8068 | -114.7533 | 1 |  | W |  | 104 |
| Running Rain | Bow | AB | 50.5045 | -114.9452 | 1 |  | Z |  | 105 |
| Silvester | Bow | AB | 50.8614 | -114.7257 | 1 |  | Z |  | 106 |
| Akolkolex | Columbia HW | BC | 50.9151 | -117.8787 | 1 |  | Z |  | 107 |
| Botts | Columbia HW | BC | 50.7986 | -116.3615 | 1 |  | Z |  | 108 |
| Cedar | Columbia HW | BC | 51.2685 | -116.9912 | 1 |  | K |  | 109 |
| Driftwood | Columbia HW | BC | 50.8700 | -116.4734 | 1 |  | Z |  | 110 |
| Ladybird | Columbia HW | BC | 49.4514 | -117.8259 | 1 |  | V |  | 111 |
| Neale | Columbia HW | BC | 51.4593 | -117.0917 | 1 |  | Z |  | 112 |
| West Fisher | Fisher | MT | 48.0537 | -115.5182 | 1 |  | Z |  | 113 |
| Doctor | Kootenai HW | BC | 50.0911 | -116.0867 | 1 |  | Z |  | 114 |
| Lodgepole | Kootenai HW | BC | 49.2819 | -114.8143 | 1 |  | X |  | 115 |
| Lussier | Kootenai HW | BC | 50.0686 | -115.5088 | 1 |  | Z |  | 116 |
| Tributary to Albert | Kootenai HW | BC | 50.5663 | -115.5596 | 1 |  | Z |  | 117 |
| Little Eightmile | Lemhi | ID | 44.7685 | -113.4424 | 1 |  | F |  | 118 |
| Reservoir | Lemhi | ID | 44.7202 | -113.1261 | 1 |  | F |  | 119 |
| Big | Lower Middle Fork Salmon | ID | 45.0732 | -115.3535 | 1 |  | Z |  | 120 |
| Flume | Lower Middle Fork Salmon | ID | 44.8181 | -114.5326 | 1 |  | F |  | 121 |
| MF Salmon | Lower Middle Fork Salmon | ID | 44.9995 | -114.7292 | 1 |  | F |  | 122 |
| NF Smith | Lower Middle Fork Salmon | ID | 45.1895 | -115.3464 | 1 |  | F |  | 123 |
| WF Cedar | Lower North Fork Clearwater | ID | 46.9419 | -115.8490 | 1 |  | T |  | 124 |
| Albert | Middle Clark Fork | MT | 46.9743 | -114.2704 | 1 |  | Z |  | 125 |
| Second | Middle Clark Fork | MT | 47.1279 | -114.7446 | 1 |  | Z |  | 126 |
| Siegel | Middle Clark Fork | MT | 47.2967 | -114.7817 | 1 |  | Z |  | 127 |
| Beaver | Middle Salmon-Panther | ID | 45.2825 | -114.2774 | 1 |  | F |  | 128 |
| Beaver | Middle Salmon-Panther | ID | 45.2837 | -114.2854 | 1 |  | F |  | 129 |
| Salmon | Middle Salmon-Panther | ID | 45.3463 | -113.9195 | 1 |  | Z |  | 130 |
| Weaver | Moyie HW | BC | 49.4034 | -116.0229 | 1 |  | Z |  | 131 |
| Harvey | NF Flathead HW | BC | 49.2491 | -114.5677 | 1 |  | Z |  | 132 |
| Lynx | Oldman | AB | 49.4961 | -114.4960 | 1 |  | Z |  | 133 |
| Oldman | Oldman | AB | 50.0570 | -114.5917 | 1 |  | Z |  | 134 |
| Twin | Oldman | AB | 50.1885 | -114.4420 | 1 |  | Z |  | 135 |
| Vicary | Oldman | AB | 49.8014 | -114.4648 | 1 |  | Z |  | 136 |
| Hunt | Priest | ID | 48.5809 | -116.7722 | 1 |  | AA |  | 137 |
| Lime | Priest | ID | 48.8966 | -116.9642 | 1 |  | Z |  | 138 |
| Bear | South Fork Salmon | ID | 44.8332 | -115.5125 | 1 |  | F |  | 139 |
| Halfway | South Fork Salmon | ID | 44.7791 | -115.5419 | 1 |  | F |  | 140 |
| NF Coeur d'Alene | Upper Coeur d'Alene | ID | 47.9586 | -116.0987 | 1 |  | V |  | 141 |
| Ross | Upper Kootenai | MT | 48.1831 | -115.9788 | 1 |  | K |  | 142 |
| WF Quartz | Upper Kootenai | MT | 48.5255 | -115.7228 | 1 |  | Z |  | 143 |
| Dagger | Upper Middle Fork Salmon | ID | 44.4676 | -115.3303 | 1 |  | F |  | 144 |
| EF Sulphur | Upper Middle Fork Salmon | ID | 44.5492 | -115.4636 | 1 |  | F |  | 145 |
| Garden | Upper Middle Fork Salmon | ID | 44.7427 | -115.1376 | 1 |  | F |  | 146 |
| MF Salmon | Upper Middle Fork Salmon | ID | 44.7304 | -115.0185 | 1 |  | F |  | 147 |
| MF Salmon | Upper Middle Fork Salmon | ID | 44.5670 | -115.3007 | 1 |  | N |  | 148 |
| Alpine | Upper Salmon | ID | 43.9046 | -114.9227 | 1 |  | F |  | 149 |
| Alpine | Upper Salmon | ID | 43.9068 | -114.9322 | 1 |  | Z |  | 150 |
| Cinnibar | Upper Salmon | ID | 44.3564 | -114.4992 | 1 |  | F |  | 151 |
| Island | Upper Salmon | ID | 44.0960 | -114.5929 | 1 |  | Y |  | 152 |
| Jordan | Upper Salmon | ID | 44.4303 | -114.7272 | 1 |  | F |  | 153 |
| Morgan | Upper Salmon | ID | 44.6517 | -113.9324 | 1 |  | F |  | 154 |
| Road | Upper Salmon | ID | 44.1569 | -114.1735 | 1 |  | F |  | 155 |
| Rough | Upper Salmon | ID | 44.2413 | -114.8006 | 1 |  | F |  | 156 |
| Salmon | Upper Salmon | ID | 44.2538 | -114.5629 | 1 |  | F |  | 157 |
| Salmon | Upper Salmon | ID | 44.5447 | -114.1767 | 1 |  | F |  | 158 |
| Salmon | Upper Salmon | ID | 44.6874 | -114.0524 | 1 |  | F |  | 159 |
| Salmon | Upper Salmon | ID | 44.4258 | -114.2225 | 1 |  | F |  | 160 |
| Salmon | Upper Salmon | ID | 44.2482 | -114.5303 | 1 |  | U |  | 161 |
| Tributary to Challis | Upper Salmon | ID | 44.5037 | -114.4571 | 1 |  | F |  | 162 |
| Magruder | Upper Selway | ID | 45.6854 | -114.7750 | 1 |  | V |  | 163 |
| Trapper | Big Hole | MT | 45.6425 | -112.8711 | 9 |  |  | x | 164 |
| Little Boulder | Bitterroot | MT | 45.7288 | -114.2451 | 10 |  |  | x | 165 |
| Overwhich | Bitterroot | MT | 45.7176 | -114.0799 | 8 |  |  | x | 166 |
| WF Bitterroot | Bitterroot | MT | 45.5040 | -114.3334 | 9 |  |  | x | 167 |
| Morrell | Blackfoot | MT | 47.2992 | -113.4639 | 8 |  |  | x | 168 |
| Nevada | Blackfoot | MT | 46.8177 | -112.5681 | 9 |  |  | x | 169 |
| Snowbank | Blackfoot | MT | 47.0561 | -112.6494 | 6 |  |  | x | 170 |
| Musselshell | Clearwater | ID | 46.3524 | -115.7558 | 7 |  |  | x | 171 |
| 4th of July | Coeur d'Alene Lake | ID | 47.6106 | -116.5178 | 10 |  |  | x | 172 |
| Big Cherry | Fisher | MT | 48.1990 | -115.5969 | 9 |  |  | x | 173 |
| Cow | Fisher | MT | 48.2257 | -115.2143 | 9 |  |  | x | 174 |
| West Fisher | Fisher | MT | 48.0427 | -115.4746 | 9 |  |  | x | 175 |
| Schwartz | Flint-Rock | MT | 46.6941 | -113.8228 | 10 |  |  | x | 176 |
| MF Little Timber | Lemhi | ID | 44.5867 | -113.4618 | 6 |  |  | x | 177 |
| Big | North Fork Flathead | MT | 48.5269 | -114.3391 | 10 |  |  | x | 178 |
| Black Bear | South Fork Flathead | MT | 47.7446 | -113.3834 | 10 |  |  | x | 179 |
| Beaver | St. Joe | ID | 47.0458 | -115.4580 | 9 |  |  | x | 180 |
| Bird | St. Joe | ID | 47.2889 | -115.5919 | 10 |  |  | x | 181 |
| Snowshoe | Upper Clark Fork | MT | 46.6860 | -112.4654 | 10 |  |  | x | 182 |
| Spotted Dog | Upper Clark Fork | MT | 46.4483 | -112.5164 | 6 |  |  | x | 183 |
| Twin Lakes | Upper Clark Fork | MT | 46.0631 | -113.2235 | 8 |  |  | x | 184 |
| Independence | Upper Coeur d'Alene | ID | 47.8806 | -116.4098 | 10 |  |  | x | 185 |
| Laverne | Upper Coeur d'Alene | ID | 47.7075 | -116.3756 | 9 |  |  | x | 186 |
| WF Eagle | Upper Coeur d'Alene | ID | 47.7367 | -115.8078 | 10 |  |  | x | 187 |
| Rail | Upper John Day | OR | 44.3462 | -118.5664 | 30 |  |  | x | 188 |
| Roberts | Upper John Day | OR | 44.3332 | -118.5781 | 22 |  |  | x | 189 |

<sup>1</sup>HU, hydrologic unit. For Canadian sites, this refers to the basin.
<sup>2</sup>*N*, sample size. Values = 1 except sites for which SNP analyses were also performed.
<sup>3</sup>MG, mitogenomic sequence. Bolded labels denote a haplotype is shared. All individuals were used in microsatellite analyses (except sample 44).
<sup>4</sup>ND2, haplotypes of NADH dehydrogenase 2.
<sup>5</sup>SNP, single nucleotide polymorphism analysis.

Whole genomic DNA was extracted from tissue samples using the QIAGEN DNeasy Tissue Kit (Qiagen, Valencia, CA, USA). Extracts were converted into indexed genomic libraries using the Illumina TruSeq DNA HT kit (version 2). To enrich samples for mitochondrial DNA targets, we developed hybridization-enrichment baits based on the complete mitochondrial genome sequence for Lahontan cutthroat trout *O. c. henshawi* (NCBI accession AY886762). Baits and blocking probes were designed and synthesized by MYcroarray, LLC (version 2; Ann Arbor, MI), and hybridization-enrichment reactions followed their protocol with a 12-cycle final amplification. Enriched libraries were pooled to 48-plex and sequenced using two lanes of 101- base-pair, single-end reads on the Illumina HiSeq2000 at the Center for Genome Research and Biocomputing at Oregon State University (http://cgrb.oregonstate.edu/). Indices were demultiplexed using CASAVA v. 1.8.2, and individual sample mitochondrial genomes were assembled using the *O. c. henshawi* reference and the CLC-Bio v. 7.0 assembler, with the following quality and alignment parameters: quality score limit of 0.05, maximum number of ambiguities per read = 2; mismatch cost = 3, indel cost = 3, indel open cost = 6, indel extend cost = 1, length and similarity fraction = 0.95, and non-specific matches mapped randomly. Individual samples were represented by an average of 3.674 million microreads (minimum = 281,755; maximum = 13,176,436); for assembly, we used a maximum of 4 million reads per sample. Short read sequences from this study are available under BioProject ID PRJNA389467 from National Center for Biotechnology Information.

In the microsatellite analysis, we conducted analyses of 10 loci used in previous studies of westslope cutthroat trout: OclMSU14, OclMSU15, OclMSU17, OclMSU21, OclMSU23, OclMSU24, OclMSU25, OclMSU27, OclMSU30, OclMSU33 (Vu and Kalinowski 2009; Drinan et al. 2011). The reaction volume (10 μl) contained 1.0 μl DNA, 1 × reaction buffer (Applied Biosystems), 2.0 mM MgCl_2_, 200 μM of each dNTP, 1 μM reverse primer, 1 μM dye-labeled forward primer, 1.5 mg/ml BSA, and 1U Taq polymerase (Applied Biosystems). The PCR profile was 94 °C/5 min, [94 °C/1 min, 55 °C/1 min, 72 °C/30 s] × 40 cycles. The resultant products were visualized on a LI-COR DNA analyzer (LI-COR Biotechnology).

For the SNP analyses, we used Competitive Allele Specific PCR (KASPar) assays (KBiosciences, Hoddesdon, Herts, England) to amplify 52 variable nuclear loci (Harwood and Phillips 2011; Kalinowski et al. 2011; Amish et al. 2012; Campbell et al. 2012; Pritchard et al. 2012). We used a suite of 44 diagnostic SNP alleles to ensure no fish introgressed with rainbow trout or Yellowstone cutthroat trout *O. c. bouvieri* alleles were included in the analysis (McKelvey et al. 2016). The PCR touchdown profile contained an initial annealing temperature of 65 °C and decreased by 0.80 °C per cycle until most cycles ran at 57 °C. We visualized PCR products on an EP1 Reader (Fluidigm) and determined individual genotypes using Fluidigm SNP Genotyping Software.

### Phylogenetic analyses

We conducted phylogenetic analyses on the three aforementioned sources of data, plus a fourth, to assess intraspecific diversity in westslope cutthroat trout.

Mitogenomic sequences were aligned in MAFFT version 7 (Katoh and Standley 2013), followed by minor manual adjustments. We used MitoAnnotator (Iwasaki et al. 2013) to order and identify gene regions. We followed Satoh et al. (2016) and the mitotRNAdb database (http://mttrna.bioinf.uni-leipzig.de/mtDataOutput/) to identify paired and unpaired regions in rRNAs and tRNAs and evaluate whether nucleotide insertions or deletions were likely to represent genotyping errors; we subsequently made 12 adjustments to the 1.6 million nucleotides in the dataset. Unique haplotypes were identified using DAMBE version 6 (Xia 2017) and used to construct neighbor-joining and maximum-likelihood phylogenetic trees. To portray evolutionary distances among and between clades, we used MEGA 7.0 (Kumar et al. 2016) to construct a neighbor-joining tree based on the MCL model with gamma-distributed evolutionary rates and some invariant positions, bootstrapped 1,000 times. We included sequences from Lahontan cutthroat trout, greenback cutthroat trout *O. c. stomias*, and rainbow trout as outgroups (GenBank accessions NC006897, KP013107, KP013117, KP013084, DQ288268-71, KP085590). Pairwise genetic distances (as a percentage) between samples and groups were based on the number of nucleotide differences. For the maximum-likelihood analysis, we used PartitionFinder 2.0 (Lanfear et al. 2016) to select the best-fitting partitioning scheme as measured by AICc, constrained to the suite of evolutionary models considered by RAxML (Stamatakis 2014) and excluding all outgroups. Data subsets included 1) codon position by gene for the protein-coding genes (39 subsets); 2) paired and unpaired regions among tRNAs and the origin of replication for the light strand (2 subsets) and between rRNAs (4 subsets); 3) intergenic spacers and the non-coding portions of the control region (1 subset); and 4) conserved sequence blocks of the control region, nucleotides coding for more than one gene, and tRNA anticodons (1 subset). Because RAxML will only consider a single evolutionary model for the entire suite of partitions, we then compared AICc scores among maximum likelihood models using the GTR and GTRGAMMA evolutionary models, and chose the model with the best score. We then ran RAxML version 8.1.21 implemented through the RAxMLGUI (Silvestro and Michalak 2012) and set for rapid bootstrapping (1,000 bootstraps) and a thorough ML search. We recognized potential units of conservation as reciprocally monophyletic clades with > 70% bootstrap support in the mitogenomic maximum-likelihood phylogeny. We did not single out subclades therein with high levels of support unless: 1) they differed from all other subclades by more than the average intra-clade divergence (0.04%) or 2) they had been previously recognized as a potential unit of conservation, i.e., fish in the Missouri River basin (Drinan et al. 2011).

Further evaluation of these clades as potential units of conservations was undertaken by analyzing a dataset combining all ND2 sequences of westslope cutthroat trout from Loxterman and Keeley (2012) and one additional specimen in GenBank (*n* = 93, accessions EU186799, JQ747580–596; Table 1) with those examined in the mitogenomic analyses (including the outgroups). Some of these specimens (*n* = 26) were used in both analyses, reducing the number of unique specimens (*n* = 163). We built a neighbor-joining tree using unpartitioned data bootstrapped 1,000 times in MEGA 7.0, for which the Tamura-Nei model with gamma-distributed rates was the best-fitting evolutionary model based on AICc scores.

Because the microsatellites and variable SNPs for westslope cutthroat trout were developed (Campbell et al. 2012) almost entirely from specimens representing the two most northeasterly and least genetically divergent clades identified in the mitogenomic analysis in the Missouri, Kootenai, and Clark Fork River basins, ascertainment bias was more likely to prevent the phylogenetic tree topology from representing the species tree (Lachance and Tishkoff 2013) but less likely to incorrectly assign individuals to groups (Bradbury et al. 2011). Consequently, we evaluated whether putative groups identified in structure 2.3.4 (Hubisz et al. 2009), using either the microsatellite or SNP datasets, matched those of the best-supported clades in the mitogenomic analyses. We applied settings recommended by Falush et al. (2003) for detecting subtle population subdivision by using the admixture model, correlated allele frequencies among populations, and an allele frequency distribution parameter (λ) set to 1. We allowed structure to infer the value of the model’s Dirichlet parameter (α) for the degree of admixture from the data. In the SNP analysis, we used *K* = 15 (the number of major river basins) as the maximum number of potential groups and completed 20 replicate runs for each value of *K*. In the microsatellite analysis, we used *K* = 10 (the number of mitogenomic clades) as the maximum number. In both analyses, we set the burn-in period to 100,000 MCMC repetitions and the postburn-in period to 100,000, ignoring the user-defined population-of-origin or sampling location for each individual (i.e., adopting non-informative priors). We attempted to identify an optimal value of *K* by calculating the maximum likelihood value and Δ*K* (Evanno et al. 2005) using the program structure harvester (Earl and vonHoldt 2012).

Stocking of genetically pure westslope cutthroat trout was inferred from the appearance of mitochondrial haplotypes outside their river basin of origin, or of individuals in nuclear analyses that assigned to groups related to those that founded the primary broodstocks in Idaho and Montana. For Montana, we regarded samples from the South Fork Flathead River (sample codes 35, 94, and 179; Table 1) as representative of broodstock genotypes. Samples from the Priest River basin (codes 24, 137, and 138) were deemed to represent fish from the Idaho broodstock, as well as the King’s Lake broodstock in Washington. The Falls Creek sample (code 60) represented Washington’s Twin Lakes broodstock, but these genotypes were not found outside the eastern Cascades river basins.

## Results

### Mitochondrial analyses

Hybridization-capture probes allowed us to enrich mitochondrial genome sequences to a very high level (mean mitochondrial representation = 57.5% of all sequences), resulting in mitochondrial genome assemblies represented by extremely high mean depth (range: 997— 20,208 reads), and complete coverage for all positions in the genome. Complete mitogenomes (16,676 nucleotides) included 13 protein-coding genes, 2 rRNAs, 22 tRNAs, and several noncoding regions (Table 2), and revealed 89 haplotypes among the 96 specimens. The mitogenomic maximum-likelihood phylogeny recovered two main lineages (mean difference between lineages based on number of nucleotide differences, 0.42%; mean difference within lineages, 0.06–0.11%) that contained nine highly supported, monophyletic clades and one subclade (Figure 2; Table 3). The topology and levels of support in the mitogenomic neighbor-joining analysis were effectively identical (results not shown). One lineage (hereafter, the southern lineage) contained six clades (1–5b) largely associated with individual river basins: the John Day River (1) in Oregon; the upper Coeur d’Alene (2) and St. Joe Rivers (3) in the Spokane River basin in Idaho; the North Fork Clearwater River (4) in Idaho; and the lower and middle Salmon River (5a) and upper Salmon River (5b) in Idaho. The latter two clades, however, overlapped geographically within the basin. The four clades (6–9) in the other main lineage (the northern lineage) were less geographically circumscribed: the Clearwater headwaters (6), which included specimens from the upper Selway and South Fork Clearwater Rivers in Idaho; the Clearwater-eastern Cascades (7), with representatives from tributaries to the Middle Fork Clearwater River in Idaho and from the Wenatchee River, Lake Chelan, and Methow River basins on the eastern side of the Cascade Range in Washington; and a neoboreal (8) clade, which included all river basins in Alberta, British Columbia, Montana, northern Idaho, and northeastern Washington. Most of these constitute basins that were covered by continental ice sheets or the proglacial lakes at their margins. Disjunct representatives of this clade, however, were found in the Salmon, Clearwater, and Spokane River basins in Idaho. Also recovered in these analyses was a well-supported subclade within the neoboreal group that represented all specimens (save one) from the Missouri River (9) basin in Montana.

**Table 2.**
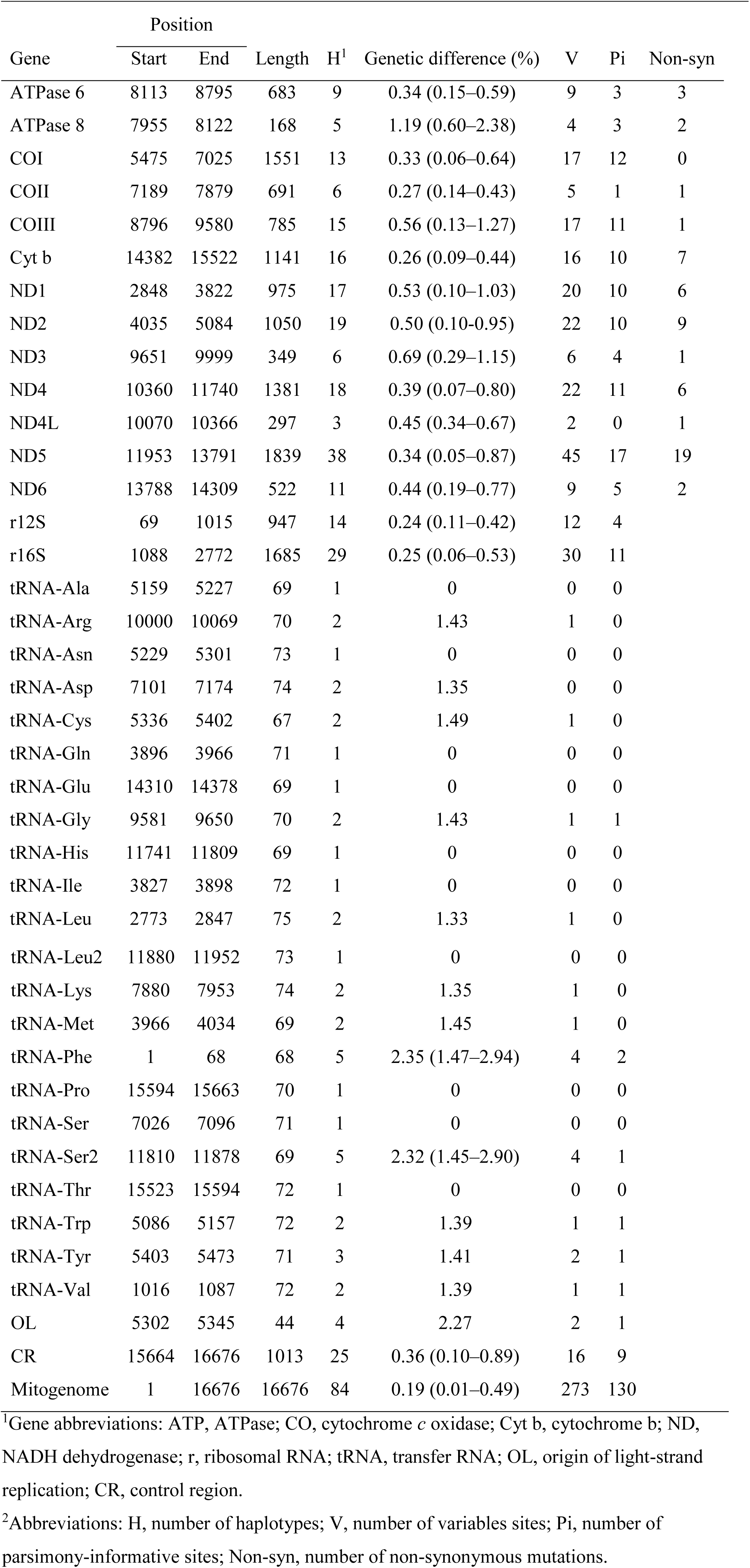
Haplotype diversity and variation by gene for 96 westslope cutthroat trout mitogenomic sequences.

| Gene | Position |  | Length | H<sup>1</sup> | Genetic difference (%) | V | Pi | Non-syn |
| --- | --- | --- | --- | --- | --- | --- | --- | --- |
|  | Start | End |  |  |  |  |  |  |
| ATPase 6 | 8113 | 8795 | 683 | 9 | 0.34 (0.15–0.59) | 9 | 3 | 3 |
| ATPase 8 | 7955 | 8122 | 168 | 5 | 1.19 (0.60–2.38) | 4 | 3 | 2 |
| COI | 5475 | 7025 | 1551 | 13 | 0.33 (0.06–0.64) | 17 | 12 | 0 |
| COII | 7189 | 7879 | 691 | 6 | 0.27 (0.14–0.43) | 5 | 1 | 1 |
| COIII | 8796 | 9580 | 785 | 15 | 0.56 (0.13–1.27) | 17 | 11 | 1 |
| Cyt b | 14382 | 15522 | 1141 | 16 | 0.26 (0.09–0.44) | 16 | 10 | 7 |
| ND1 | 2848 | 3822 | 975 | 17 | 0.53 (0.10–1.03) | 20 | 10 | 6 |
| ND2 | 4035 | 5084 | 1050 | 19 | 0.50 (0.10–0.95) | 22 | 10 | 9 |
| ND3 | 9651 | 9999 | 349 | 6 | 0.69 (0.29–1.15) | 6 | 4 | 1 |
| ND4 | 10360 | 11740 | 1381 | 18 | 0.39 (0.07–0.80) | 22 | 11 | 6 |
| ND4L | 10070 | 10366 | 297 | 3 | 0.45 (0.34–0.67) | 2 | 0 | 1 |
| ND5 | 11953 | 13791 | 1839 | 38 | 0.34 (0.05–0.87) | 45 | 17 | 19 |
| ND6 | 13788 | 14309 | 522 | 11 | 0.44 (0.19–0.77) | 9 | 5 | 2 |
| r12S | 69 | 1015 | 947 | 14 | 0.24 (0.11–0.42) | 12 | 4 |  |
| r16S | 1088 | 2772 | 1685 | 29 | 0.25 (0.06–0.53) | 30 | 11 |  |
| tRNA-Ala | 5159 | 5227 | 69 | 1 | 0 | 0 | 0 |  |
| tRNA-Arg | 10000 | 10069 | 70 | 2 | 1.43 | 1 | 0 |  |
| tRNA-Asn | 5229 | 5301 | 73 | 1 | 0 | 0 | 0 |  |
| tRNA-Asp | 7101 | 7174 | 74 | 2 | 1.35 | 0 | 0 |  |
| tRNA-Cys | 5336 | 5402 | 67 | 2 | 1.49 | 1 | 0 |  |
| tRNA-Gln | 3896 | 3966 | 71 | 1 | 0 | 0 | 0 |  |
| tRNA-Glu | 14310 | 14378 | 69 | 1 | 0 | 0 | 0 |  |
| tRNA-Gly | 9581 | 9650 | 70 | 2 | 1.43 | 1 | 1 |  |
| tRNA-His | 11741 | 11809 | 69 | 1 | 0 | 0 | 0 |  |
| tRNA-Ile | 3827 | 3898 | 72 | 1 | 0 | 0 | 0 |  |
| tRNA-Leu | 2773 | 2847 | 75 | 2 | 1.33 | 1 | 0 |  |

|  |  |  |  |  |  |  |  |
| --- | --- | --- | --- | --- | --- | --- | --- |
| tRNA-Leu2 | 11880 | 11952 | 73 | 1 | 0 | 0 | 0 |
| tRNA-Lys | 7880 | 7953 | 74 | 2 | 1.35 | 1 | 0 |
| tRNA-Met | 3966 | 4034 | 69 | 2 | 1.45 | 1 | 0 |
| tRNA-Phe | 1 | 68 | 68 | 5 | 2.35 (1.47–2.94) | 4 | 2 |
| tRNA-Pro | 15594 | 15663 | 70 | 1 | 0 | 0 | 0 |
| tRNA-Ser | 7026 | 7096 | 71 | 1 | 0 | 0 | 0 |
| tRNA-Ser2 | 11810 | 11878 | 69 | 5 | 2.32 (1.45–2.90) | 4 | 1 |
| tRNA-Thr | 15523 | 15594 | 72 | 1 | 0 | 0 | 0 |
| tRNA-Trp | 5086 | 5157 | 72 | 2 | 1.39 | 1 | 1 |
| tRNA-Tyr | 5403 | 5473 | 71 | 3 | 1.41 | 2 | 1 |
| tRNA-Val | 1016 | 1087 | 72 | 2 | 1.39 | 1 | 1 |
| OL | 5302 | 5345 | 44 | 4 | 2.27 | 2 | 1 |
| CR | 15664 | 16676 | 1013 | 25 | 0.36 (0.10–0.89) | 16 | 9 |
| Mitogenome | 1 | 16676 | 16676 | 84 | 0.19 (0.01–0.49) | 273 | 130 |

<sup>1</sup>Gene abbreviations: ATP, ATPase; CO, cytochrome *c* oxidase; Cyt b, cytochrome b; ND, NADH dehydrogenase; r, ribosomal RNA; tRNA, transfer RNA; OL, origin of light-strand replication; CR, control region.
<sup>2</sup>Abbreviations: H, number of haplotypes; V, number of variables sites; Pi, number of parsimony-informative sites; Non-syn, number of non-synonymous mutations.

**Figure 2.**
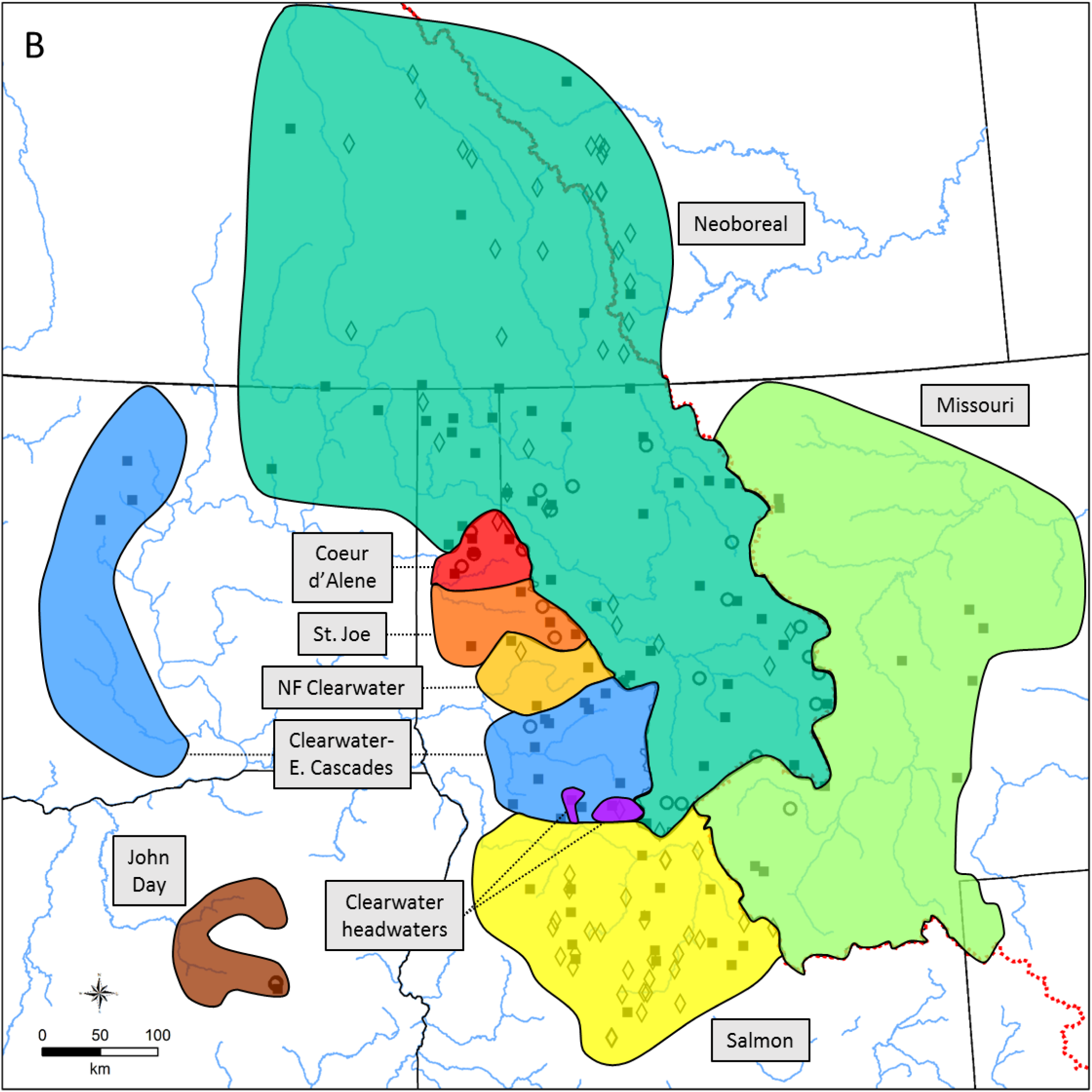
Maximum-likelihood phylogeny of individual mitogenomic haplotypes (*n* = 89) of westslope cutthroat trout from across its historical range. Support (as a percentage, based on 1,000 bootstraps) is given above each node. Only support > 70% is shown. Mitogenomic clades discussed in the text are identified by a unique color. Sample labels are in Table 1.

The analysis of ND2 sequences from fish genotyped for this study and from public databases had 27 haplotypes, including 8 not observed in the mitogenomic analysis (Table 1). Consistent with the previous analysis, the ND2 neighbor-joining tree resolved the two main lineages and recovered similar geographic structure (Figures 3), but with less resolution and support of additional clades despite greater geographic coverage. This analysis diagnosed the John Day, Coeur d’Alene, and St. Joe River clades, and grouped but did not distinguish among the North Fork Clearwater and Salmon River specimens. It also pooled all Clearwater River and eastern Cascade specimens, and did not resolve Missouri River specimens as a distinct group within the neoboreal clade.

**Figure 3.**
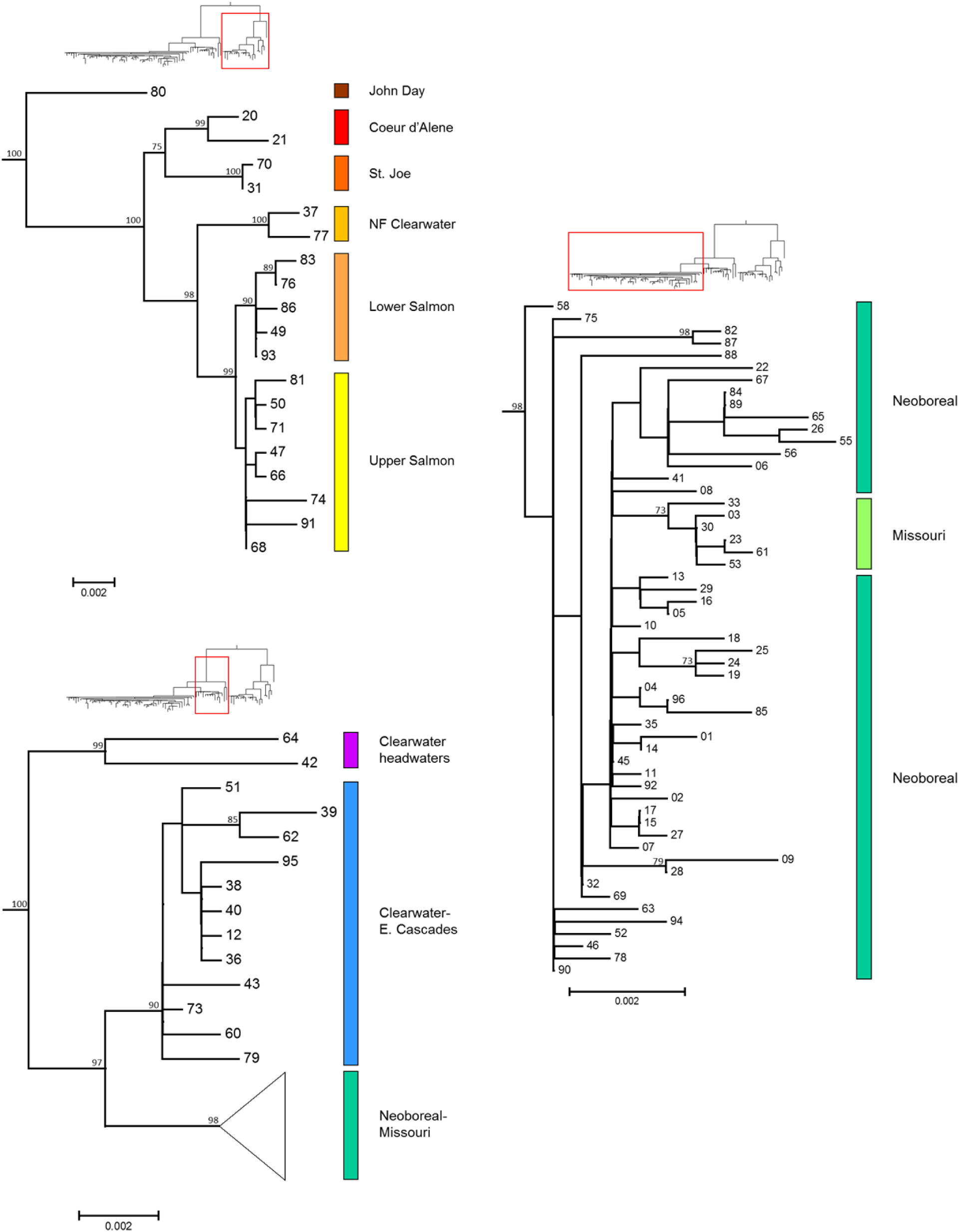
Neighbor-joining phylogeny of individual ND2 haplotypes (*n* = 27) of westslope cutthroat trout from across its historical range. Support (as a percentage, based on 1,000 bootstraps) is given above each node. Only support > 70% is shown. Mitogenomic clades discussed in the text are identified by a unique color. ND2 haplotype labels are in Table 1.

### Nuclear analyses

In the structure analysis of SNPs, measures of the optimal number of groups were not concordant. The log-likelihood estimate peaked at the maximum number of groups (*K* = 15) whereas Δ*K* identified *K* = 2 as the optimum number of clusters. Given this outcome, we examined how assignment to groups changed across the entire range of values (Figure 4 depicts *K* = 4, 8, and 15). The intermediate value (*K* = 8) was equivalent to the number of mitogenomic clades geographically represented in the SNP analysis (samples equivalent to the North Fork Clearwater River and Clearwater River headwater samples were not available). Although these results broadly corroborated the mitogenomic classification, there were important differences. Specimens from the John Day River were the first group to be segregated as distinct (at *K* = 3). Specimens from the Salmon River formed a distinct group at intermediate and higher levels of *K*, as did those from the Missouri River. The Clearwater River and eastern Cascades clade grouped at the lower levels of *K*, but split into distinct clusters at the highest levels of resolution. More complex were patterns among the Coeur d’Alene River, St. Joe River, and neoboreal clades. Specimens from the Coeur d’Alene and St. Joe Rivers largely assigned to a single group at all levels of *K*, but this group also included specimens from the neoboreal clade from the lower Clark Fork, Pend Oreille, and lower Kootenai River basins for lower levels of *K*. The rest of the neoboreal clade tended to split along major watershed boundaries at higher levels of *K*, particularly the upper Kootenai River.

**Figure 4.**
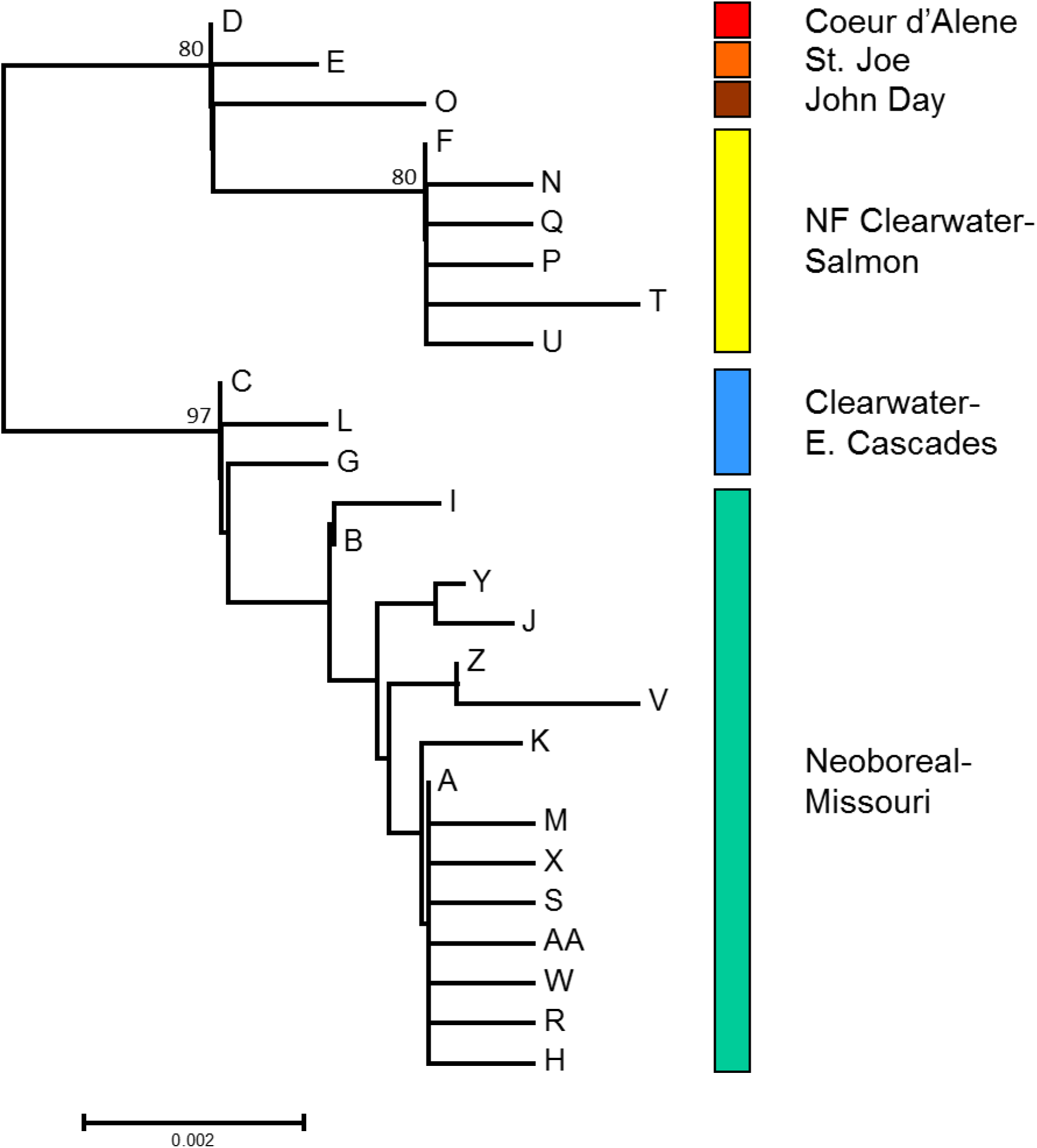
Assignment of westslope cutthroat trout (*n* = 524) to *K* = 4, 8, or 15 genetic clusters analyzed at 52 variable SNP loci using structure. Each individual (denoted by a single bar) is represented by the proportion of their genotype that was assigned to each of *K* clusters. Samples are sorted geographically by basin, but structure was run with no geographic priors. The left vertical axis identifies major drainage basins of each sample and the right vertical axis the mitogenomic clades. Clades for the North Fork Clearwater River and Clearwater River headwaters were not included because no fish from these areas were examined in the SNP analysis.

In the microsatellite analyses, *K* = 5 (Figure 5) had the largest log-likelihood estimate and the third-highest Δ*K* (which peaked at *K* = 2). The smaller sample sizes in this analysis afforded less resolution but nonetheless highlighted groups consistent with those identified previously: 1) the John Day and Clearwater Rivers and eastern Cascades basins; 2) the Salmon River; 3) the Coeur d’Alene and St. Joe Rivers, some samples from the Pend Oreille and lower Kootenai Rivers, one from downstream on the Columbia River, and one from the Fraser River; 4) the Clark Fork and upper Kootenai Rivers in the Columbia basin and the Oldman and Bow Rivers in the South Saskatchewan basin; and 5) the Missouri River.

**Figure 5.**
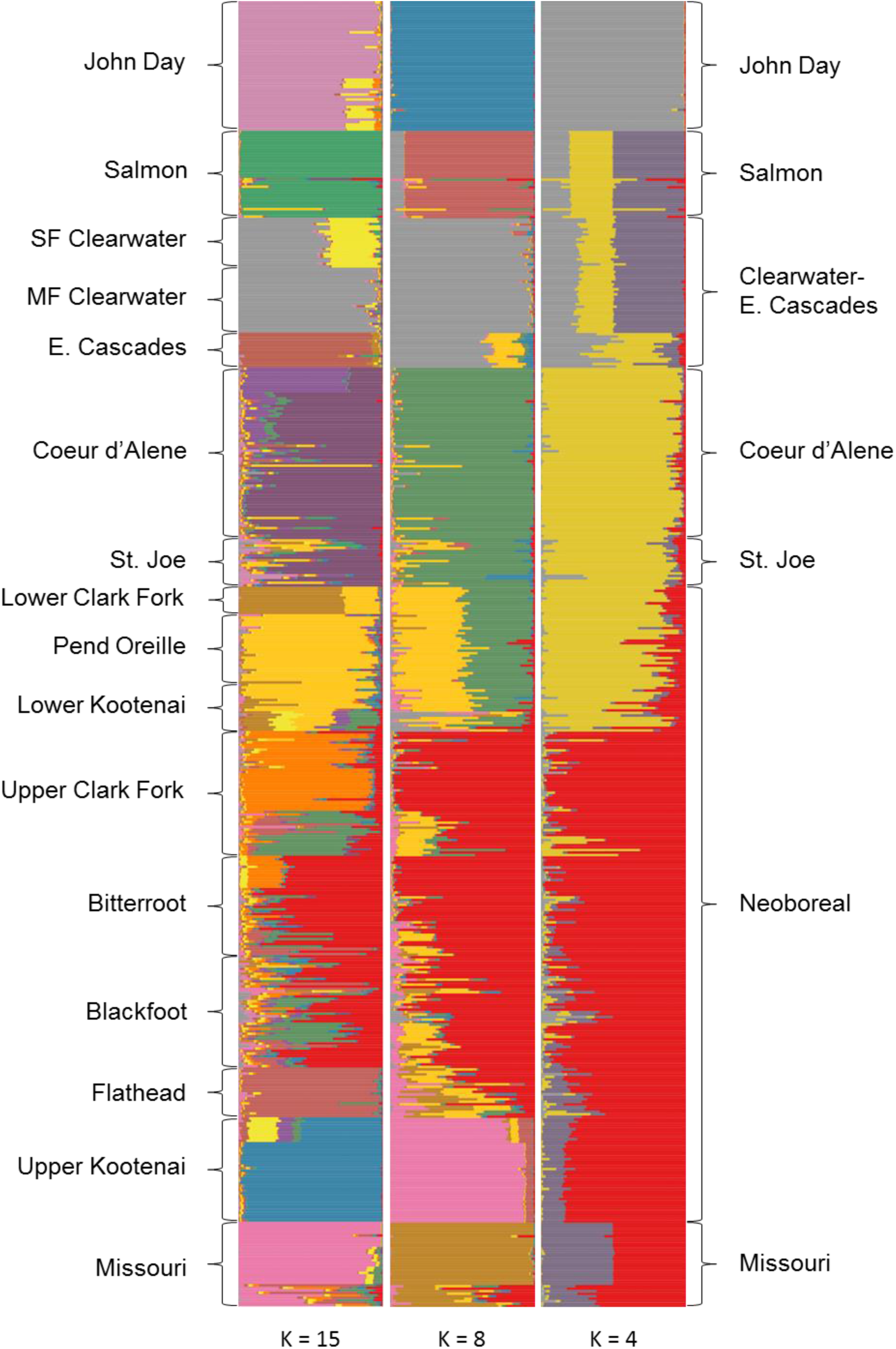
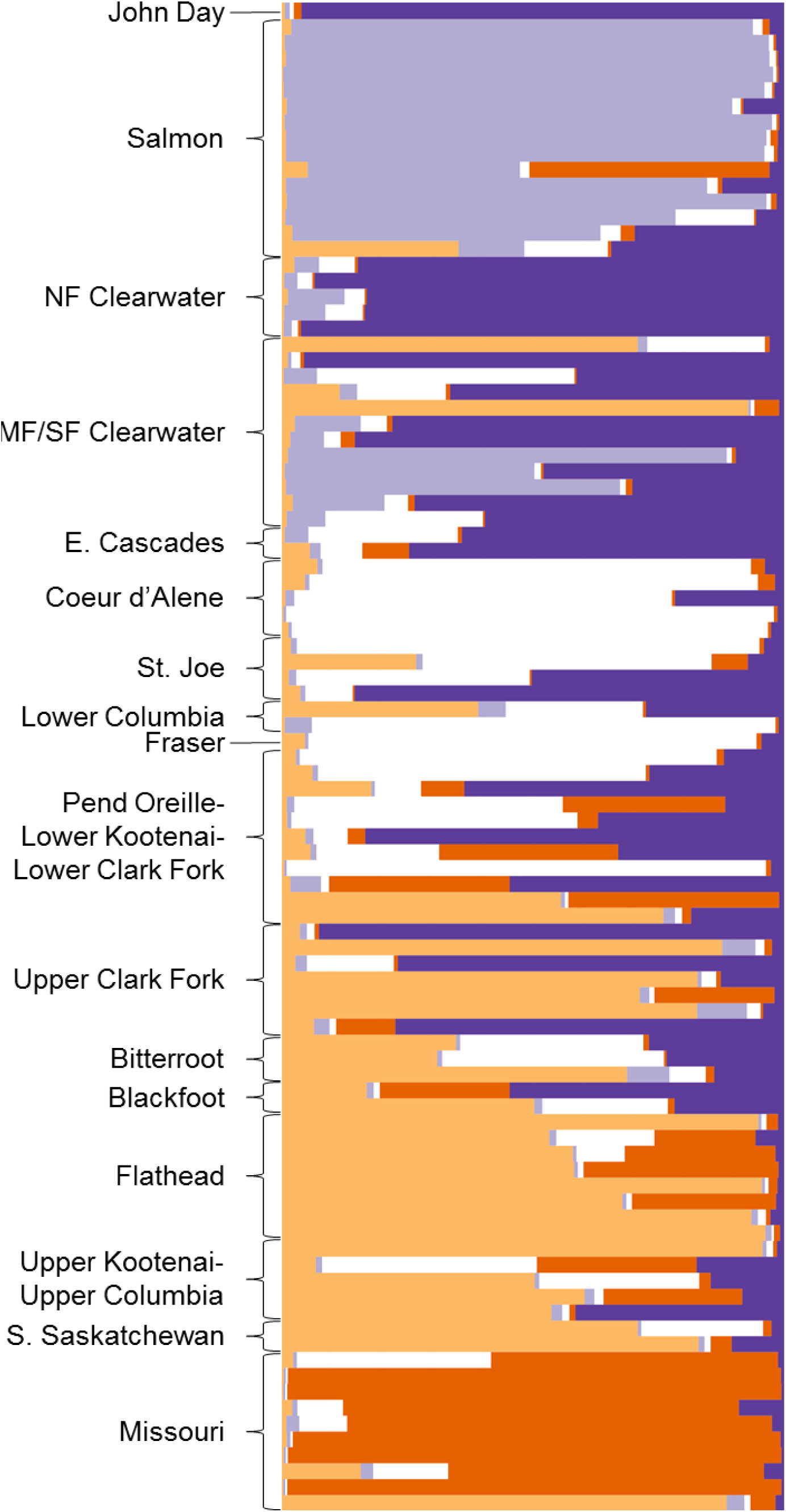
Assignment of westslope cutthroat trout (*n* = 95) to *K* = 5 genetic clusters analyzed at 10 microsatellite loci using structure. Each individual (denoted by a single bar) is represented by the proportion of their genotype that was assigned to each of *K* clusters. Samples are sorted geographically by basin, but structure was run with no geographic priors. The left vertical axis identifies major drainage basins of each sample.

### Stocked fish

The genotypes of one or more fish at as many as 18 sites were suspected of being influenced by introductions of nonintrogressed but nonindigenous westslope cutthroat trout (Table 4). The bulk of these observations were from Idaho, and involved the appearance of mitochondrial haplotypes typical of the neoboreal group in the Salmon and Clearwater River basins. Although we regarded specimens with neoboreal haplotypes in the St. Joe and Coeur d’Alene River basins, primarily from the lowermost portions of both basins, as being nonindigenous, this interpretation in uncertain. Fish with either the neoboreal or southern haplotypes tended to share a nuclear genotype that was also typical of fish found farther north in the Pend Oreille, lower Clark Fork, lower Kootenai, middle Columbia, and Fraser River basins. Thus, the St. Joe or Coeur d’Alene River specimens may have been indigenous representatives of fish that colonized basins to the north, including that from which Idaho’s broodstock was derived, rather than reflect recent introductions of that hatchery stock. In Montana, the two likely examples of nonindigenous specimens were from disparate parts of the Missouri River basin that harbored fish having mitochondrial or nuclear genotypes akin to those from west of the Continental Divide.

**Table 3.** Mean pairwise genetic divergence (%) among clades (below the diagonal) and within clades (on the diagonal).

|  | John Day | Coeur<br>d'Alene | St. Joe | North Fork<br>Clearwater | Salmon<br>A | Salmon<br>B | Clearwater<br>headwaters | Clearwater-E. Cascades | Neoboreal | Missouri |
| --- | --- | --- | --- | --- | --- | --- | --- | --- | --- | --- |
| John Day | NA |  |  |  |  |  |  |  |  |  |
| Coeur d'Alene | 0.20 | 0.05 |  |  |  |  |  |  |  |  |
| St. Joe | 0.19 | 0.10 | 0.01 |  |  |  |  |  |  |  |
| North Fork<br>Clearwater | 0.24 | 0.16 | 0.16 | 0.04 |  |  |  |  |  |  |
| Salmon A | 0.21 | 0.13 | 0.14 | 0.11 | 0.02 |  |  |  |  |  |
| Salmon B | 0.21 | 0.13 | 0.14 | 0.11 | 0.05 | 0.03 |  |  |  |  |
| Clearwater<br>headwaters | 0.36 | 0.41 | 0.40 | 0.44 | 0.42 | 0.42 | 0.11 |  |  |  |
| Clearwater-E. Cascades | 0.34 | 0.39 | 0.39 | 0.43 | 0.40 | 0.40 | 0.14 | 0.04 |  |  |
| Neoboreal | 0.36 | 0.42 | 0.41 | 0.46 | 0.43 | 0.43 | 0.16 | 0.10 | 0.04 |  |
| Missouri | 0.37 | 0.43 | 0.42 | 0.46 | 0.43 | 0.44 | 0.17 | 0.10 | 0.04 | 0.01 |

**Table 4.** Stocking records associated with sample sites exhibiting geographically incongruent genotypes. Data are ND2 sequences, mitogenomic (MG) sequences, or SNPs. Stocking information is from the online databases for Idaho (https://idfg.idaho.gov/fish/stocking) and Montana (http://fwp.mt.gov/fishing/mFish/).

| Site code | Site | State | Data | Expected genotype | Nearby stocking | Date | Number |
| --- | --- | --- | --- | --- | --- | --- | --- |
| 25 | Mokins | ID | MG | Coeur d'Alene | Hayden Lake | 1969–2007 | 4,853,000 |
| 26 | NF St. Joe | ID | MG | St. Joe | St. Joe River | 1968–1973 | 235,000 |
| 29 | Dry Wolf | MT | MG | Missouri | No data |  |  |
| 32 | Tributary of Armstrong | ID | MG | Coeur d'Alene | Coeur d'Alene Lake | 1967–1969 | 73,000 |
| 41 | Bad Luck | ID | MG | Clearwater headwaters | Canyon Creek Lakes, White Cap Creek Lakes | 1968–2008 | 83,000 |
| 48 | Morse<sup>2</sup> | ID | SNP | Salmon | No data |  |  |
| 55 | Leiberg | ID | MG | Coeur d'Alene | Coeur d'Alene River | 1968–1972 | 153,000 |
| 59 | Yoosa<sup>1</sup> | ID | MG | Clearwater-E. Cascades | No data |  |  |
| 65 | EF Emerald | ID | MG | St. Joe | No data |  |  |
| 89 | Twin | ID | MG | Salmon B | Twin Creek, WF Bear Creek | 1990–2008 | 1,500 |
| 120 | Big | ID | ND2 | NF Clearwater-Salmon | Belvidere Lakes | 1971–1989 | 26,000 |
| 130 | Salmon | ID | ND2 | NF Clearwater-Salmon | No data |  |  |
| 141 | NF Coeur d'Alene | ID | ND2 | Coeur d'Alene | LNF Coeur d'Alene River, Spruce Creek | 1968–1972 | 264,000 |
| 150 | Alpine | ID | ND2 | NF Clearwater-Salmon | Alpine Creek Lakes | 1970–2009 | 44,000 |
| 152 | Island | ID | ND2 | NF Clearwater-Salmon | Island Lake | 1969–1999 | 14,000 |
| 163 | Magruder | ID | ND2 | Clearwater-E. Cascades | Gold Pan Lake | 1968–1994 | 18,000 |
| 164 | Trapper | MT | SNP | Missouri | No data |  |  |
| 177 | MF Little Timber | ID | SNP | Salmon | NF Little Timber Lakes | 1983–1986 | 500 |

<sup>1</sup>All fish in the SNP analysis clustered as expected.
<sup>2</sup>The fish in the mitogenomic analysis clustered as expected.

## Discussion

The phylogeographic structure of westslope cutthroat trout is consistent with unglaciated basins serving as refugia during glacial cycles, followed by extensive colonization from some of those refugia during interglacial intervals (Bernatchez and Wilson 1998; Hampe and Petit 2005). The deepest divergence among populations involved the John Day, Salmon, Clearwater, and Spokane River basins, which were south of the maximum advance of the Cordilleran ice sheet. Much lower levels of divergence characterized areas formerly occupied by the ice sheets or the glacial lakes at their margins, collectively represented by the neoboreal mitogenomic clade (and including the Missouri River basin). Although we have foregone estimating dates of divergence among clades, much of the diversity seems to be associated with recent climatic events. The radiation following the Last Glacial Maximum likely reflects the divergence within most mitogenomic clades (mean, 0.04%), whereas previous interglacials over 100 ka ago may have led to divergence between the Coeur d’Alene and St. Joe River clades, the neoboreal and the Clearwater-eastern Cascade clades, and the North Fork Clearwater and Salmon River clades, and still older events to the split between the southern and northern major lineages. Continental glaciation during previous glacial maxima is likely to have extirpated groups that advanced northward during earlier interglacial intervals. That all samples from the previously glaciated area form a single mitogenomic clade suggests that no individuals representing earlier colonizers are present in this area. Below, we consider the phylogeographic relevance of these river basins to westslope cutthroat trout in more detail.

The clade in the John Day River basin, though clearly a member of the southern major lineage, exhibited the greatest divergence from all other groups. This watershed constitutes the downstreammost extent of westslope cutthroat trout in the Columbia River basin and is separated from the next nearest populations in the eastern Cascade Range in Washington by hundreds of kilometers, and by hundreds more from the nearest members of the southern lineage in the Clearwater and Salmon Rivers in Idaho. There has been uncertainty with regard to whether these fish were indigenous or introduced from Montana, Idaho, or Washington (Gunckel 2002), but the evidence indicates that these fish constitute a distinct, relictual population of westslope cutthroat trout that did not arrive via glacial outwash floods from upstream. Whether their distribution extends to the North Fork John Day River basin, which also supports westslope cutthroat trout but for which there are extensive records of stocking of this taxon (Gunckel 2002), has not been evaluated. As alluded to earlier, the absence of westslope cutthroat trout from nearby basins in southeastern Washington and northeastern Oregon with abundant suitable habitat is somewhat surprising. That westslope cutthroat trout may have been present, however, is hinted at by the presence of their mitochondrial haplotype among anadromous rainbow trout in at least one area, the Tucannon River in Washington (Brown et al. 2004), although unrecorded stocking of westslope cutthroat trout is a plausible alternative explanation.

The Salmon River basin constitutes a modern stronghold for westslope cutthroat trout (Shepard et al. 2005). It also hosts two relatively divergent mitogenomic clades of the southern lineage which are found exclusively in this basin. Nevertheless, the clades are geographically interspersed and showed little differentiation in the nuclear analyses. Their origin may have reflected isolation in different parts of the Salmon River basin during a previous glacial interval followed by subsequent mixing. Their nearest relatives appear to be fish found in the North Fork Clearwater River to the north, but whether these groups represent descendants of one another or of simultaneous divergence in these different locations is uncertain.

The Clearwater River basin in Idaho constitutes a biodiversity hotspot for westslope cutthroat trout. It hosts a geographically complex mixture of members of both primary mitogenomic lineages and four of the 10 mitogenomic clades of westslope cutthroat trout, two of which are endemic (and a third may be introduced). Its position beyond the southerly extent of the Cordilleran ice sheet, combined with its relatively low elevation and high precipitation, seems to have contributed to the basin being a refugium for disjunct or locally endemic populations of an array of plant and animal taxa during the Last Glacial Maximum (summarized in Shafer et al. 2010). That does not suggest that all portions of the basin were amenable to westslope cutthroat trout throughout this or earlier glacial cycles. Pollen core records from near the headwaters indicate a cold, arid climate with vegetation dominated by sagebrush, and that western red cedar *Thuja plicata,* currently a dominant forest overstory species typical of mesic, montane environments, did not colonize the basin until the last 5 ka (Herring and Gavin 2015). Moreover, mountain glaciers would have been prevalent in the higher ranges, constraining westslope cutthroat trout to the lower portions of many watersheds until conditions permitted them to colonize these areas, perhaps from multiple locations within the basin.

A remarkably close relationship between the populations in the eastern Cascade Range in Washington and those in Clearwater River basin—especially the Lochsa River basin—in Idaho is supported in all analyses. This enigmatic pattern raises questions about how westslope cutthroat trout crossed the majority of the lower Snake and middle Columbia River basins in recent geological time and in which direction. The simplest explanation would be stocking of the Twin Lakes broodstock from Washington in the Clearwater River in Idaho, but there are no records to support this (Evan Brown, Idaho Department of Fish and Game, personal communication). There is, however, evidence of the disjunct distribution of undescribed species of sculpins in central Idaho basins and along the eastern Cascade Range (M. Young, unpublished data). Given that the modern distribution of these sculpins does not include lacustrine or large, warm riverine environments, it is more likely that they, and thus westslope cutthroat trout, were transported downstream from central Idaho to their present distribution in central Washington. The limited divergence in westslope cutthroat trout haplotypes between these locations also indicates that this transfer was recent, and may have involved the emptying of Lake Bonneville or the last of the Glacial Lake Missoula outburst floods.

The major forks of the Spokane River basin are the Coeur d’Alene and St. Joe Rivers. In their headwaters, these basins harbor distinct clades that are nonetheless sister to one another and represent the southern lineage of westslope cutthroat trout. A comparable pattern of relatedness and divergence was evident in the cedar sculpin *Cottus schitsuumsh*, a recently described species that is largely confined to the Spokane River basin (LeMoine et al. 2014). The lower portions of both the Coeur d’Alene and St. Joe Rivers, however, also hosted members of the mitogenomic neoboreal clade. Fish from throughout both basins shared a nuclear genotype that also appeared in fish farther north in the middle Columbia, Pend Oreille, lower Kootenai, lower Clark Fork River, and Fraser River basins, all presently unconnected to the Spokane River basin. The pattern could have arisen from 1) stocking of fish from the Priest River basin, the source of Idaho’s broodstock, throughout the upper Spokane River basin, for which there is some evidence (Table 4); or 2) ephemeral connections associated with fluctuating levels of proglacial lakes that promoted northerly expansion of genotypes from the Spokane River basin. As noted earlier, Glacial Lake Columbia reached tens of kilometers up the Coeur d’Alene and St. Joe River valleys and would have facilitated the movement of fish between these basins. This would also have provided access to the Pend Oreille basin across the low divide in Rathdrum Prairie in northern Idaho. Much of this watershed was subjected to the extreme conditions associated with the repeated, catastrophic drainage of Glacial Lake Missoula, but Glacial Lake Columbia persisted for some time after the last of these floods (Balbas et al. 2017). As the Purcell trench lobe of the Cordilleran ice sheet receded, it would have exposed the lower Kootenai River valley but blocked drainage to the north, diverting streams from that area through the trench cut by the Purcell lobe and connecting the lower Kootenai and Pend Oreille basins (Langer et al. 2011). Another connection between these basins appears to have existed at the low divide between the Bull River (lower Clark Fork) and Bull Lake (lower Kootenai) in Montana (Langer et al. 2011). Each of these connections would have provided access to fish originating from the Spokane River basin, which may have culminated in their reaching the Fraser River basin via the succession of glacial lakes trailing the receding ice sheet. That a fish from that basin was assigned to a microsatellite group representing fish from the upper Spokane River basin argues for such a connection.

Yet it is also clear that these fish—or this particular wave of colonizers—did not found the modern populations of westslope cutthroat trout inhabiting the Kootenai River above Kootenai Falls or the Clark Fork River above present-day Noxon Dam or Thompson Falls. All these fish constitute a single mitogenomic clade with a broadly shared nuclear genotype that suggests a limited number of founding individuals or a limited number of sources. Whatever their origin, this is now the most widely distributed clade of westslope cutthroat trout, and one that also displays substantial internal structure. For example, the nuclear data (e.g., SNP structure plots for *K* = 2–6) indicate historical connections between populations in upper Kootenai and Flathead River basins, for which there are a number of plausible pathways that might have existed depending on the timing of recession of particular lobes of the Cordilleran ice sheet, the presence or absence of terminal or lateral moraines that affected drainage patterns, or the existence of multiple outlets of lakes near drainage divides, e.g., between the Little Bitterroot (Flathead), Big Thompson (Clark Fork), and Fisher (upper Kootenai) Rivers, or the Stillwater (Flathead) and Tobacco (upper Kootenai) Rivers.

There is also evidence that these fish not only colonized two additional major river basins across the Continental Divide—the Missouri and South Saskatchewan Rivers—but that these appear to be independent events. Based on our results and on earlier studies (Drinan et al. 2011), the Missouri River clade is a recent derivative of the widely distributed neoboreal clade, likely founded from a single source; all individuals share two diagnostic mutations in the mitogenome (one each in tRNA for tyrosine and the control region). The nuclear data imply that their closest relatives are those in the upper Flathead River basin. A likely route is via the Middle Fork Flathead River and Nyack Creek to Summit Creek, thence to the Two Medicine River basin and the rest of the Missouri River basin. This invasion was likely to have coincided with the lingering presence of glacial lakes formed by the Laurentide ice sheet because the Two Medicine River is tributary to the Marias River, which presently flows into the Missouri River well downstream of the Great Falls of the Missouri. Because the falls constitute a barrier to upstream fish migration, this location was likely inundated by a glacial lake (e.g., Glacial Lake Great Falls had a highstand of 1200 m [Calhoun 1906], about 250 m higher than the river elevation at the falls) which would have enabled westslope cutthroat trout to access the Missouri River headwaters. Furthermore, the lack of genetic divergence (mean distance 0.01%, with several geographically distant pairs of samples with identical mitogenomic haplotypes) among westslope cutthroat trout in this basin also indicates that their recent arrival may be their first. The upper Missouri River basin was not covered by the Laurentide Ice Sheet during any portion of the Pleistocene, thus suitable habitat for cutthroat trout should have persisted throughout much of this period and permitted greater divergence. That westslope cutthroat trout did not extend their distribution to the Yellowstone River basin, where Yellowstone cutthroat trout are found, points to the absence of a migration corridor via the lower Missouri and Yellowstone Rivers for either subspecies when they colonized. That a single clade of sculpins *Cottus* sp. (Young et al. 2013; M. Young, unpublished data) and of mountain whitefish *Prosopium williamsoni* (Whiteley et al. 2006) co-occur with both subspecies in these areas, however, demonstrates that interbasin exchanges were possible at an earlier time.

The westslope cutthroat trout of Alberta in the South Saskatchewan River basin appear to have arrived via a different route. They share mitogenomic affinities with fish of the Clark Fork River basin upstream from Thompson Falls and lack the diagnostic mutations seen in the fish from the Missouri River basin. The microsatellite analyses also group these fish with those in the upper Clark Fork River, not with those from the Missouri River. That the invasions of the South Saskatchewan and upper Missouri River basins were independent events is also supported by the distribution of bull trout *Salvelinus confluentus*, a cold-water salmonid with similar habitat preferences, which are present in the former and absent from the latter (Ardren et al. 2011). Moreover, westslope cutthroat trout may have been constrained to a limited portion of the South Saskatchewan River basin. Their current distribution includes only the Bow and Oldman River basins (Fisheries and Oceans Canada 2014), and Marnell (1988) argued that they were historically absent from Waterton Lakes and the Belly River and crossed the Continental Divide farther north. Based on topography, a plausible route would have been via the Kootenay River to Summit Lake in British Columbia, thence to Crowsnest Lake in Alberta, which is part of the Oldman River basin.

Based on our combined analyses, we find evidence for nine units of conservation within this taxon (Table 5). We acknowledge that different interpretations of the importance of modern-day isolation, administrative boundaries, ecoregions, or geographic mixing of genotypes (Mee et al. 2015) could alter this total. For example, a reasoned case can be made that fish representing the neoboreal clade in the lower Kootenai, lower Clark Fork, Pend Oreille, middle Columbia, and Fraser River basins should be classified separately from those found elsewhere because their nuclear genotypes are distinct. Similarly, one could contend that fish in the eastern Cascades in Washington are distinct from those in the Clearwater River basin in Idaho based on geographic isolation and different administrative authorities. Such arguments are valid, thus we consider our estimate an initial hypothesis to be revised following additional sampling and more comprehensive analyses, but would be cautious about divisions at much smaller scales. Leary et al. (1988) reported that the bulk of genetic variation in westslope cutthroat trout in Montana was partitioned among individual populations, and consequently advocated for saving as many populations as possible to preserve the genetic legacy of this taxon. The substantial structuring of headwater populations has been observed elsewhere among westslope cutthroat trout (Taylor et al. 2003; Young et al. 2004) and other salmonid species (Ryman 1983; Wenburg and Bentzen 2001; Pritchard et al. 2009), though it may be an artifact of the widespread anthropogenic isolation of headwater trout populations (Campbell et al., this volume). In contrast, our data (also see Drinan et al. 2011) representing nearly the entire range of westslope cutthroat trout demonstrates that the primary divisions within this taxon arise at broader spatial scales, roughly conforming to major watershed boundaries and cycles of Pleistocene glaciation, which is consistent with hypotheses about rangewide genetic structure in temperate zone species (Hampe and Petit 2005). Although we concur with the intent to save as many populations as possible, the larger groups we identified are more likely to satisfy criteria for recognizing units of conservation (Waples 1991; Mee et al. 2015).

**Table 5.** Proposed units of conservation for westslope cutthroat trout.

| Name | Location | Evidence<sup>1</sup> |
| --- | --- | --- |
| John Day | Upper portion of the John Day River basin | G, MG, MS, SNP |
| Coeur d'Alene | Upper portion of the Coeur d'Alene River basin | G, MG |
| St. Joe | Upper portion of the St. Joe River basin | G, MG |
| North Fork Clearwater | Upper portion of the North Fork Clearwater River | G, MG, ND2 (in part) |
| Salmon | Salmon River basin | G, MG, ND2 (in part), MS, SNP |
| Clearwater headwaters | Headwaters of the Selway and South Fork Clearwater Rivers | G, MG |
| Clearwater-eastern Cascades | Lochsa River, lower Selway River, and lower portions of tributaries to the Middle Fork Clearwater River, and the Wenatchee River, Lake Chelan, and Methow River basins | G (in part), MG, MS, SNP |
| Neoboreal | Columbia River basin from the Sanpoil River upstream (excluding the Spokane River), Fraser River basin, and the South Saskatchewan River basin | G, MG, ND2 (in part), MS, SNP |
| Missouri | Missouri River basin | G, MG, MS, SNP |

<sup>1</sup>Evidence is from geographic discreteness (G), mitogenomic sequences (MG), microsatellites (MS), or single nucleotide polymorphisms (SNP).

Our data and that of others confirms that the westslope cutthroat trout constitutes a coherent taxon that may warrant recognition as a full species. The level of divergence within westslope cutthroat trout (0.39% in a 640-bp fragment of cytochrome c oxidase subunit I (COI), the barcode region) is comparable to that of other species of fishes (0.36%) worldwide (Ward 2009), whereas its divergence (1.25% in the COI fragment, 1.11% across the mitogenome) from its nearest relative (Lahontan cutthroat trout) attains or exceeds that of many species pairs (April et al. 2011). Nonetheless, substantial uncertainties remain with regard to the placement of westslope cutthroat trout within the overall cutthroat trout lineage. Our mitogenomic analyses affirmed that westslope cutthroat trout haplotypes are highly divergent from those of rainbow trout (4.80% across the mitogenome), and more similar to Lahontan cutthroat trout than to greenback cutthroat trout (1.58%), but there is little consensus among the different genomic regions. Others have suggested that westslope cutthroat trout may be most closely related to rainbow trout based on variation in isozyme alleles (Leary et al. 1987), to coastal cutthroat trout based on karyotypes or morphology (Loudenslager and Gall 1980; Behnke 1992), to Lahontan cutthroat trout based on mitochondrial gene sequences (Wilson and Turner 2009; Loxterman and Keeley 2012), either of the latter two based on analyses of Y chromosomes (Brunelli et al. 2013), and to all three based on a panel of variable nuclear SNP markers (Houston et al. 2012). That each analysis supports a different relationship may be more indicative of the relative strengths and weaknesses of different markers than of an ambiguous phylogeny. For example, allozymes can grossly and unpredictably underestimate haplotype variation (Buth 1984). The karyotype analysis was conducted on a sample of five westslope cutthroat trout from one hatchery broodstock (Loudenslager and Gall 1980) and its results contradict those of an earlier study with specimens from elsewhere (Simon and Dollar 1963). Both studies could be correct because geographical variation in chromosomal characteristics or multiple karyotypes within populations are common (Thorgaard 1983). Mitochondrial sequences form the basis of a multitude of projects to identify animal species (Ratnasingham and Herbert 2013), but introgression and reticulate evolution among lineages weaken relationships based solely on mitochondrial data (Taylor and Harris 2012). And all genetic analyses are vulnerable to incomplete geographic sampling, and the geographic origin and ancestry of specimens used to develop nuclear markers and diagnostic phenotypes can undermine phylogenetic inferences because of ascertainment bias (Lachance and Tishkoff 2013) or environmentally driven selection. Given the relative youth of cutthroat trout lineages, more intensive and representative sampling of the nuclear genome and of populations likely to be indigenous will be required to resolve this issue.

Although our work represents the most comprehensive effort to describe the phylogeography of westslope cutthroat trout, many details have yet to be resolved. Locations represented by one or two individuals would benefit from more thorough sampling to reveal the distribution and mixing of presumptive haplotypes and the discovery of new ones, such as in the John Day River (and its presumably stocked North Fork) in Oregon, the eastern Cascade River basins in Washington, and many portions of the Clearwater River in Idaho. Westslope cutthroat trout in the Yakima River basin are of unknown provenance, whether related to lineages in the John Day, the eastern Cascades (as implied in Figure 1), or to one as yet undescribed. Similarly, additional work to delineate hatchery stocks using nuclear markers would clarify the extent to which introgression of introduced forms affects indigenous populations, and would help avoid the use of introduced forms when attempting to establish basin-specific stocks. Despite these uncertainties, the phylogeographic patterns we have observed are concordant with the varying connections among river basins in the intermountain West throughout the late Pleistocene. These patterns can be used as a foundation to understand phylogeographic structure in other freshwater taxa, albeit with an awareness of the nuances associated with the long evolutionary history of cutthroat trout in western North America.

## Acknowledgements

We thank M. Small of the Washington Department of Fish and Wildlife and J. Neal of the Oregon Department of Fish and Wildlife for contributing samples. T. Cross of the Rocky Mountain Research Station assisted with the structure analysis of SNPs, and D. Horan helped prepare Figure 1. Comments by Kurt Fausch and two anonymous reviewers markedly improved the manuscript. This work was funded by the RIM Board of the U.S. Forest Service’s Region 1, the Rocky Mountain Research Station, and the Pacific Northwest Research Station.

